# Splenic clearance of rigid erythrocytes as an inherited mechanism for splenomegaly and natural resistance to malaria

**DOI:** 10.1101/2022.03.21.485136

**Authors:** Benoît Henry, Geoffroy Volle, Hilaire Akpovi, Laure Gineau, Camille Roussel, Papa Alioune Ndour, Félicien Tossou, Felipe Suarez, Friso Palstra, Aurélie Fricot, Charlotte Chambrion, Julien Solinc, Julie Nguyen, Mathilde Garé, Florentin Aussenac, Charles-Henry Cottart, Christine Keyser, Rafiou Adamou, Magali Tichit, David Hardy, Nadine Fievet, Jérôme Clain, André Garcia, David Courtin, Olivier Hermine, Audrey Sabbagh, Pierre Buffet

**Affiliations:** Université de Paris, Biologie Intégrée du Globule Rouge, UMR_S1134, BIGR, INSERM, Paris, France; Laboratoire d’Excellence Gr-Ex, Paris, France; Institut National de la Transfusion Sanguine, Paris, France; Service des maladies infectieuses et tropicales, APHP, Hôpital Necker Enfants Malades, Centre d’Infectiologie Necker-Pasteur, Institut Imagine, Paris, France; Service des maladies infectieuses et tropicales, APHP.Université Paris Saclay, Hôpital Bicêtre, Le Kremlin-Bicêtre, France; CERPAGE (Centre d’Etude et de Recherche sur les Pathologies Associées à la Grossesse et à l’Enfance), Cotonou, Bénin; Université de Paris, IRD, MERIT, Paris, France; Centre Interfacultaire de Formation et de Recherche en Environnement pour le Développement Durable (CIFRED), Université d’Abomey-Calavi, Cotonou, Bénin; Ministère de la Santé, Cotonou, Bénin; Service d’hématologie adultes, APHP, Hôpital Necker Enfants Malades, Paris, France; Université de Paris, INSERM U1163, CNRS ERL 8654, Paris, France; Service de biochimie générale, APHP, Hôpital Necker Enfants Malades, Faculté de pharmacie, Paris, France; Université de Paris, CNRS, UMR 8045, Paris, France.; Institut Pasteur, Experimental Neuropathology Unit, Paris, France; Institut Pasteur, Paris, France

**Keywords:** malaria, *falciparum*, spleen, erythrocytes, ethnic groups, heritability, splenomegaly, genome-wide association study

## Abstract

In malaria-endemic areas, subjects from specific groups like Fulani have a peculiar protection against malaria, with high levels of IgM but also frequent anemia and splenomegaly. The mechanisms underlying this phenotype remain elusive. In Benin, West Africa, we measured the deformability of circulating erythrocytes in genetically distinct groups (including Fulani) living in sympatry, using ektacytometry and microsphiltration, a mimic of how the spleen clears rigid erythrocytes. Compared to non-Fulani, Fulani displayed a higher deformability of circulating erythrocytes, pointing to an enhanced clearance of rigid erythrocytes by the spleen. This phenotype was observed in individuals displaying markers of *Plasmodium falciparum* infection. The heritability of this new trait was high, with a strong multigenic component. Five of the top 10 genes selected by a population structure-adjusted GWAS, expressed in the spleen, are potentially involved in splenic clearance of erythrocytes (*CHERP*, *MB*, *PALLD*, *SPARC*, *PDE10A*), through control of vascular tone, collagen synthesis and macrophage activity. In specific ethnic groups, genetically-controlled processes likely enhance the innate retention of infected and uninfected erythrocytes in the spleen, explaining splenomegaly, anemia, cryptic intrasplenic parasite loads, hyper-IgM, and partial protection against malaria. Beyond malaria-related phenotypes, inherited splenic hyper-filtration of erythrocytes may impact the pathogenesis of other hematologic diseases.

**Research in context:** *Evidence before this study:* The genetic background of individuals influences their susceptibility to infectious diseases. Specific human groups, like the Fulani in Africa, react to malaria parasites (named *Plasmodium*) in a specific way. Upon infection, Fulani develop a grossly enlarged spleen, and high levels of anti-*Plasmodium* antibodies in their blood. They also carry smaller numbers of parasites in their blood, and thus are considered partially protected against malaria. The mechanisms underlying this natural protection, different from other natural protective mechanisms such as the sickle cell trait, are not well understood. Malaria impairs the deformability of red blood cells and the spleen is a key organ to controlling red blood cell quality. We have recently demonstrated that red blood cells containing live malaria parasites accumulate intensely in the spleen of subjects with long term exposure to these parasites. Enhanced retention of infected and uninfected red blood cells in the spleen would explain why the spleen is larger and why lower numbers of parasites are left in circulation. We thus explored whether the retention of infected and uninfected red blood cells could explain why Fulani are partially protected against malaria. Because it is unethical to perform spleen puncture or biopsies for research purposes, our explorations were indirect by carefully analyzing the properties of circulating red blood cells in a large number of subjects and by assessing whether observations could be explained by their genetic make-up.

*Added value of this study:* In more than 500 subjects, we confirmed the high frequency of large spleens in Fulani and, through 2 different methods, we demonstrated an enhanced deformability of their circulating red blood cells, that likely stems from the more efficient removal of the less deformable ones. This enhanced deformability was found to be inheritable based on carefully collected family links and refined analysis of genetic markers.

*Implications of all the available evidence:* Our findings indicate that genes potentially driving the filtration of red blood cells by the spleen likely influence how subjects in specific groups in Africa and elsewhere react to malaria. While most previous hypotheses pointed to conventional immunological mechanisms as the trigger, we propose that a simple physiological mechanism that controls the quality of red blood cells may drive natural protection from malaria even before the intervention of immunological cells. A better understanding of these processes is of great importance in the context of malaria elimination efforts. These findings may also have an impact on the understanding of other red blood cell-related disorders, such as inherited red cell diseases, in which splenic filtration of abnormal red blood cells may precipitate splenic complications.

## Introduction

Malaria due to *Plasmodium falciparum* is the most frequent acquired red blood cell (RBC) disease worldwide. Natural protection against malaria has been extensively studied, predominantly leading to the identification of RBC polymorphisms such as sickle cell trait as protective genotypes [1]. Fulani, traditional nomadic pastoralists living in West Africa, display a specific phenotype in relation to malaria. Compared to other ethnic groups living in sympatry, Fulani have a higher prevalence of splenomegaly [2–7], higher total IgM levels [8], and a stronger humoral immune response against plasmodial antigens [9, 10]. In some but not all studies, malaria-related fever and parasitemia are lower and/or less prevalent in Fulani than in other groups [4,6,11–15], and their hemoglobin levels are generally lower than in control groups [16]. Not least, the immune response against *P. falciparum* in Fulani displays specificities with regards to cytokines production [17, 18], monocytes activity [19], B-cell subpopulations [20], and regulatory T-cell function [21].

Intra-erythrocytic stages of *P. falciparum* are mostly confined to the blood [22] and splenic circulations [23, 24]. The mechanical properties of *P. falciparum*-infected RBC have been extensively studied over the last 40 years. A decrease in deformability of RBC infected with mature, but also ring-stage parasites, has been observed and contributes to vascular flow obstruction observed in severe malaria [25, 26]. This reduced deformability extends to uninfected RBC [27] and is correlated with disease severity [28]. The human spleen plays a key role in the quality control of the circulating RBC. The splenic red pulp senses the surface and mechanical properties of RBC in the specific anatomic structures of the cords and inter-endothelial slits. After observing the mechanical retention of parasitized RBC in human spleens perfused ex-vivo [29], we had proposed that in malaria-infected subjects, the clearance of infected (and possibly uninfected) RBC by the spleen could modulate the size of the parasite biomass as well as where the antigenic stimulus predominates. A stringent retention of RBC in the spleen would confer protection by reducing the parasite biomass in circulation, but would also be pathogenic by precipitating splenomegaly and anemia [30]. Recent observations of tissue samples from splenectomy in a malaria-endemic area have revealed the intense accumulation of RBC containing live malaria parasites in the human spleen [23, 24], bringing further substance to our hypothesis. Following this trail, we have quantified the deformability of circulating RBC in a multi-ethnic cohort of Beninese subjects living in sympatry in an area of high malarial transmission, using a filtration method that mimics the mechanical sensing of RBC by the human spleen [31]. The objectives were to deliver a more precise definition of malaria-related phenotypes in African groups with regards to RBC and spleen, and to investigate the potential genetic components underlying these phenotypes.

## Methods

### Study setting and clinical evaluation

This study was part of a program led by the French Research Institute for Development (Institut de Recherche pour le Développement, IRD) and Université de Paris, aiming at deciphering the genetic, parasitological, environmental and behavioural determinants of the relative protection of Fulani from malaria (BAObAB study: Biocultural AdaptatiOn to malaria in Atacora, northern Benin). This study recruited nuclear families and, in some instances, extended families.

The study was conducted in 4 rural villages (Goufanrou, Gorgoba, Tamande, Kouboro) located 40 km from the Natitingou city center, 3-15 km apart, in the Birni district of the Atacora department, a region of northwestern Benin with one of the highest prevalence of malaria in the country [32].

Malaria transmission is seasonal and *Plasmodium falciparum* is the predominant parasite species transmitted by *Anopheles gambiae s.s.* (85 %) and *Anopheles arabiensis* (15 %) [33]. Highest infection rates occur during the rainy season, from May to late October, with mean sporozoite inoculation rates reaching two infective bites per person per night (A. Djènontin, personal communication, September, 2018), and then drop during the dry season from November to April [34]. Most of the field work and experiments were performed at the end of the transmission season in December, 2017. This cross-sectional survey systematically recorded clinical and biological information related to malaria (i.e., temperature, spleen enlargement rate, hemoglobin level, parasite carriage and parasite density). Included subjects underwent a physical examination, including tympanic temperature measurement, spleen palpation, signs of portal hypertension in case of splenomegaly. A rapid hemoglobin determination using the Hemocue© (Radiometer), and a rapid diagnostic test for malaria (RDT; SD BIOLINE Malaria Ag P.f©, Abbott) using immunochromatographic detection of Histidine Rich Protein 2 (HRP2) of *P. falciparum* were also performed. A thick smear was prepared from all subjects, fixed with methanol, stained with Giemsa and read at x1000 magnification. Parasitemia was determined by counting parasites against 500 leucocytes and assuming a total leucocyte count of 8000/mm3. A thick blood smear was declared negative when no parasite was detected in 200 fields. Only *P. falciparum* mono infections were analyzed. Treatment of malaria (artemether-lumefantrine, as recommended by the Beninese National Malaria Control Program) was provided to the study population at no cost.

Individual information (age, gender, ethnic background, symptoms, recent use of antimalarial agents) was collected. Ethnicity was assessed through self-declaration, and, to be eligible for the study, participants must had identified their parents and four grandparents as belonging to the same ethnic group as themselves (Bariba, Otamari, Gando or Fulani). Self-declarations were controlled and confirmed by community health workers and community leaders from each village. Familial relationships were built by questioning each of the household members and were further confirmed by village authorities. In the case of conflicting information, the ambiguous kinship relationships were clarified by genotyping individuals with the GlobalFiler® kit (ThermoFischer Scientific). Reconstruction of family relationships resulted in 116 distinct pedigrees, including 103 nuclear and 13 extended families. Among the 420 individuals included in the heritability and familial correlation analyses, we identified 337 parent-offspring pairs, 138 sibling pairs, 59 half-sibling pairs, 16 grandparent-grandchild pairs, 47 avuncular pairs, 54 cousin pairs, 16 half-avuncular pairs, 6 great avuncular pairs and 4 half-cousin pairs. A field survey revealed that the 116 studied families lived in 189 distinct households at the time of study. Close members of a same pedigree did not always live in the same house and thus did not share the same household environment. Conversely, there were many cases where unrelated individuals or distant relatives (higher than the first degree) shared the same house.

### Blood samples

Processing of blood samples is shown in Figure 4. Venous peripheral blood (4-6 ml for adults, 1- 3 mL for children and infants) was collected on-site on EDTA (BD vacutainer©) and immediately stored at 4-8°C until use. After sedimentation, plasma was collected and cryopreserved at -20°C, then -80°C and shipped to the central laboratory on dry ice. RBC were transferred into SAGM (saline, adenine, glucose, mannitol, Macopharma©) a preservation medium used for transfusion, at 5% haematocrit, and shipped to the central laboratory at 4°C for phenotypic analysis. We had previously shown the stability of measures for up to 3 weeks under these storage conditions (Supplementary Figure 4a and 4b).

### Total IgM measurement

Immunoturbidimetric measurement was performed on an Architec C1600 analyzer (Abbott Laboratories Diagnostics Division) according to the manufacturer recommendations (Abbott technical sheet: 1E01-21).

### Circulating RBC deformability assays

Ektacytometry [35] was performed on SAGM-preserved RBC, diluted in viscous iso-osmolar polyvinylpyrrolidone (PVP; RR Mechatronics©) at 0·5% haematocrit, temperature 37°C, between 9 and 17 days post sampling. We used a Laser-assisted Optical Rotational Cell Analyzer (RR Mechatronics©), which evaluates the diffraction pattern of RBC through a laser, when elongated between two rotating cylinders at different shear stresses between 0·3 and 30 Pascals (Pa). Results were expressed as an Elongation Index (EI), which is the ratio of the difference between the 2 axes of the ellipsoidal diffraction pattern and the sum of these 2 axes. We collected EI at low (0·3 Pa, 3 Pa), intermediate (1·69 Pa), and high shear stress (30 Pa), the latter being close from what is encountered in the splenic red pulp microcirculation [36].

Microsphiltration (“microsphere filtration”; MS) assesses the ability of RBC to squeeze through narrow slits between metallic microbeads. This technique reflects the filtration of normal and abnormal RBC in the splenic red pulp, as assessed by direct comparison with retention rates in isolated-perfused human spleens [31]. It has been adapted to 96 and 384-wells microplates, allowing us to process numerous blood samples over a short period of time [37]. Microsphiltration was performed using frozen (-20°C) 96-wells plates thawed just before use. Plates were prepared with a Tecan® automat as described [37]. Control RBC were obtained from an O-negative healthy blood donor (Etablissement Français du Sang, Lille, France), and were carboxyfluorescein diacetate succinyl ester (CFSE)-stained (20 μmol/l; Life Technologies). For each sample tested, the cell mix consisted of 95% control stained RBC, and 5% tested RBC (left unlabelled) from Beninese subjects. The hematocrit of the mix was 1%, and cells were diluted in Krebs-Henseleit buffer (Sigma-Aldrich) modified with 2·1 g/l of sodium bicarbonate and 0·175 g/l of calcium chloride, and supplemented with Albumax© (0·5%; Life Technologies), at a pH of 7·4. Each RBC sample was tested in triplicate. Results were expressed as retention or enrichment rates (RER), using flow cytometry analysis of upstream and downstream samples (BD FacsCantoII©, BD Biosciences). RER was calculated as follows: ([proportion of unlabelled RBC in downstream sample] minus [proportion of unlabelled RBC in upstream sample])/proportion of unlabelled RBC in upstream sample. Positive values, called enrichment, indicate that RBC from the subject are more deformable than control RBC, whereas negative values, called retention, indicate the opposite. Positive controls were glutaraldehyde (0·8%)-treated RBC, which are completely retained; negative controls were unstained control RBC. For each subject, the mean value of RER was calculated after exclusion of outliers, and results were normalized to the negative control value. Experiments were performed 15 to 17 days after RBC collection. Cytometric data were acquired with FACSDIVA software v6.1.3 (BD Biosciences) and analyzed with FlowJo v10.0 software (FlowJo, LLC).

### Nucleic acid extraction and real-time quantitative PCR (qPCR)

Total nucleic acids were extracted from filter papers containing approximately 10 μL of dried whole blood using an optimized Chelex protocol. Final eluate volume was 140 μL. Two species- specific qPCRs were performed. The first qPCR targeted a single-copy gene (human albumin or alb; forward and reverse primers were GCTGTCATCTCTTGTGGGCTG and AAACTCATGGGAGCTGCTGGTT, 140-bp amplicon [38]) and served as an internal extraction control. The second qPCR targeted a multiple copy *P. falciparum* gene (cytochrome b or pfcytb, with an average 15-20 copies per parasite; forward and reverse primers were TTGGTGCTAGAGATTATTCTGTTCCT and GGAGCTGTAATCATAATGTGTTCGTC, 188-bp amplicon). qPCRs were carried out in 15 μl in a 384-well plate using SensiFASTTM SYBR Hi-ROX kit (Bioline), 0·5 μM of each forward and reverse primer and 2·5 μl of template DNA. Amplifications were performed in an Applied-Biosystems 7900 HT Fast Real Time PCR machine (Thermo Fisher Scientific): 95°C for 3 min; then 40 cycles of 95°C for 15 sec, and 63°C for 1 min. Several controls were run on each 384-well plate: water, *Plasmodium*-free human DNA, and seven DNA from *Plasmodium*-infected blood isolates (*P. vivax*, *P. ovale*, *P. malariae*, and four different *P. falciparum*). For tested DNA samples, the alb and pfcytb qPCRs were performed once and as duplicates, respectively, on each plate. Tested DNA samples were scored as *P. falciparum*-positive when two conditions were met: 1) the alb qPCR gave a threshold value (Ct) < 28 with a melt curve and temperature similar to corresponding positive controls; and 2) the two pfcytb qPCR replicates gave a Ct ≤ 38 with a melt curve and temperature similar to corresponding positive controls. All amplification and melt curves were individually inspected by eye.

### Definitions used for phenotypic analysis

Anemia was defined according to WHO criteria [39], based on the age-related hemoglobin levels as follows: lower than 11 g/dL in 0-4-year-olds; lower than 11·5 g/dL in 5-11-year-olds; lower than 12 g/dL in 12-14-year-olds; lower than 12 g/dL in in adult women and lower than 13 g/dL in adult men. Fever was defined as a tympanic temperature of 38°C or more.

### Heritability analysis

Heritability (*h*^2^) was estimated using the variance-components approach implemented in the SOLAR package, which simultaneously utilizes data on all family relationships [40]. This method applies maximum likelihood estimation to a mixed effects model that incorporates fixed effects for known covariates and variance components for genetic and environmental effects. The genetic component is assumed to be polygenic with no variation attributable to dominance components. A variance component was included in the model to represent the possible shared environmental effect related to the household (environmental exposures shared by individuals living in the same house at the time of study), also referred to as ‘household effects’ (*c*^2^). Heritability was estimated as the ratio of additive genetic variance to the total phenotypic variance unexplained by covariates. Estimates of the means and variances of components of the models were obtained by maximum likelihood methods and significance was determined by likelihood ratio tests. Intrafamilial correlation coefficients were estimated using the pairwise weighting scheme for all available pairs of relatives in the pedigrees using the FCOR program within the SAGE v6.3 software package (S.A.G.E. [2016], Release 6.4: http://darwin.cwru.edu). Correlations were calculated for the residual trait values obtained from a multivariate linear regression model, including relevant covariates (here RDT positivity and ethnicity). Homogeneity testing of correlations among subtypes (e.g. mother-offspring and father-offspring) within main type (e.g. parent-offspring) was also performed. Under the null hypothesis of homogeneity, the test statistic has an approximate χ2 distribution with degrees of freedom equal to the number of subtypes minus one.

### Genotyping and data quality control

Genotyping was conducted at the Centre National de Recherche en Génomique Humaine (CNRGH, CEA, Evry, France). Before genotyping, a quality control was systematically performed on each DNA sample, as detailed in Supplementary Methods.

After quality control, DNA samples were aliquoted in 96-well plates (JANUS liquid handling robot, Perkin Elmer) for genotyping; sample tracking was ensured by a systematic barcode scanning for each sample. Genotyping was performed on Illumina HumanOmni5-4v1 chips, in accordance with the standard protocol of Illumina Infinium HD assay, as detailed in the Supplementary Methods.

Standard quality control metrics were then applied to the genome-wide data of the 347 genotyped samples using PLINK v1.90b4.6 software [41]. Single nucleotide polymorphisms (SNPs) with a call-rate < 0·98 (n=96,834), a minor allele frequency (MAF) < 0·01 (n=1,740,582) or displaying significant departure from Hardy-Weinberg equilibrium (HWE) (*p* < 10^−4^) (n=1,714) were removed from the dataset, as well as duplicated (n=70,807) and non-autosomal SNPs (n=109,008). For DNA samples, individuals with discordant sex information (n =16), an autosomal heterozygosity rate more than three standard deviations on either side of the mean (n=4), and sample duplicates (n=4) were excluded. All remaining samples had a call rate >0·98 (mean = 0·998) and were kept for analysis. These quality control checks resulted in a final set of 2,281,899 high-quality SNPs genotyped in 323 individuals.

### Genotype imputation

Genome-wide imputation was performed using the IMPUTE2 v2.3.2 algorithm [42]. First, genotypes were prephased into best-guess haplotypes with SHAPEIT v2 to increase the computational efficiency of downstream steps [43–45]. We used a window size of 2 Mb, an effective population size of 17,469 (as recommended by the software providers for African populations) and the duoHMM method that allows to incorporate the known pedigree information to improve phasing [46]. Then, IMPUTE2 was run on the prephased haplotypes using the cosmopolitan panel of reference haplotypes from the 1000 Genomes Project Phase 3 (October 2014 release; *n* = 2,504) [47]. The genome was split up into non-overlapping segments of 5 Mb. We used imputation parameter settings of k =500, Ne= 20,000 and an internal buffer region of 250 kb. Imputation was performed in parallel for each segment, and segments were reconstructed into chromosomes once all imputations had finished. Stringent post-imputation quality and accuracy filters were applied to genotypes resulting from the imputation (info ≥ 0·8, certainty ≥ 0·9 and concordance ≥ 0·9 between directly measured and imputed genotypes after masking input genotypes). Further, we applied the same SNP QC criteria as the aforementioned genotyped analysis (removal of variants with call-rate < 0·98, HWE test p < 10−4 or MAF < 0·01). Following imputation and quality control, 14,350,106 genotyped or imputed SNPs and indels were available for the genome-wide association analysis.

### Genome-wide association study

The genome-wide association analysis was performed in R (version 2.15.1; http://CRAN.R-project.org) using the GRAMMAR–GC (Genome-wide Rapid Association using Mixed Model and Regression – Genomic Control) method [48] implemented in the GenABEL software (version 1.8-0) [49]. This method is suitable for the analysis of data in which both ethnic admixture and close family relationships are present while successfully controlling the overall genomic inflation factor to an appropriate level to avoid false-positive results. Phenotypes are regressed on genotypes one SNP at a time in GRAMMAR-GC in three steps. First, phenotypes were corrected by accounting for population stratification and familial dependence among individuals by using the genomic kinship matrix (pairwise kinship coefficients amongst sampled individuals estimated on the basis of genome-wide SNP data). These correlation-free residuals were then used as dependent quantitative traits for association analysis of each SNP using a linear regression model. Finally, genomic control (GC) was applied to correct the test statistic using the genomic inflation factor (λ), which is the regression coefficient of the observed statistic on the expected statistic. We used a genome-wide significance threshold of 5 × 10^-8^. Additional loci with a p-value ≤ 10^-6^ were listed as suggestive. The quantile–quantile (QQ) plot of the test statistic was visually inspected and the genomic inflation factor λ was calculated to evaluate potential inflation of the test statistic due to sample relatedness and population structure.

### Gene mapping and biological prioritization

Loci that showed evidence of association at p≤5 x 10^-6^ with the deformability of circulating RBC (as assessed by microsphiltration, after adjustment for relevant covariates) were mapped to genes using the SNP2GENE tool implemented in FUMA (http://fuma.ctglab.nl/ [50]), a web plateform designed for post-processing of GWAS results and prioritizing of genes. FUMA annotates candidate SNPs in genomic risk loci and subsequently maps them to prioritized genes based on (i) physical position mapping on the genome, (ii) expression quantitative trait loci (eQTL) mapping, and (iii) 3D chromatin interactions (chromatin interaction mapping). It incorporates the most recent bioinformatics databases, such as Combined Annotation-Dependent Depletion score (CADD score [51]), regulatory elements in the intergenic region of the genome (RegulomeDB [52]), Genotype-Tissue Expression (GTEx), and 3D chromatin interactions from Hi–C experiments [53].

We considered as genomic risk loci those with at least one SNP (lead SNP) with a p-value of association ≤ 10^-6^. In the initial step, SNPs in LD (r^2^ > 0·1, estimated from the African reference panel of KGP phase 3) in a 250 kb window were considered as belonging to the same risk locus. Thereafter, a set of candidate SNPs were selected, based on LD (r^2^ >0·6) with the lead SNP (defined as SNP with the lowest p value) from either our data set or from the African population data of KGP phase 3. If two independent significant lead SNPs (r² < 0·1) were close to each other (significant loci of interest less than 250 kb apart), the two regions were merged into a single significant region. The major histocompatibility complex (MHC) region was excluded from analysis.

Positional mapping was performed using maximum distance from SNPs to gene of 10 kb and on the basis of annotations obtained from ANNOVAR. Criteria to define deleterious SNPs were a CADD score >12·37 which indicates potential pathogenicity or an RDB score ≤ 4 which indicates that the SNP likely lies in a functional location (categories 1 and 2 of RegulomeDB classification identifed as “likely to affect binding”).

EQTL mapping was performed using eQTL data from tissue types considered as relevant for our phenotype of interest. The selection included Alasoo 2018 macrophage; BLUEPRINT monocyte, neutrophil and T-cell; CEDAR B-cell CD19, monocyte CD14, neutrophil CD15, T-cell CD4, T- cell CD8, and platelet; Fairfax 2012 B-cell CD19; Fairfax 2014 IFN24; GENCORD T-cell; Kasella 2017 T-cell CD4 and T-cell CD8; Lepik 2017 blood; Naranbhai 2015 neutrophil CD16; Nedelec 2016 macrophage; Quach 2016 monocyte; TwinsUK blood; van der Wijst scRNA eQTLs (9 tissues); DICE (15 tissues); Westra et al (2013) Blood eQTL Browser; Zhernakova et al (2017) BIOS QTL Browser; eQTLGen (cis and trans eQTLs); GTEx (Genotype-Tissue Expression) v8 blood, blood vessel and spleen tissues. Only significant SNP-gene pairs (with a false discovery rate (FDR) ≤ 0.05) were selected.

Chromatin interaction mapping was performed from Hi–C experiments data of three tissues identifed as relevant for our phenotype of interest: aorta, spleen and mesenchymal stem cell (GSE87112), as well as from the FANTOM EP correlations cell type and organ databases. Candidate genes were selected whenever their promoter region (250bp upstream and 500bp downstream of the TSS) belonged to a locus significantly interacting with candidate lead SNPs or SNPs in LD with them in the selected tissues. Additional filtering was performed to restrict the candidate genes (i) to those with a promoter region associated with SNPs overlapping predicted enhancer regions in selected epigenomes and (ii) to those having promoter regions overlapping predicted promoter regions in selected epigenomes. Filtering was restricted to specific epigenomes: blood (23 distinct tissues), spleen and aorta (E065). FDR was used to correct for multiple testing (FDR < 1.10^-6^ for chromatin interaction, the default value in FUMA).

### Functional annotation and prediction of SNPs and genes

Annotation of SNPs associated with RBC deformability was conducted using FUMA and the Variant Annotation Integrator tool [54]. The selected lead SNP from each associated locus was annotated using a deleteriousness score (CADD score), a score assessing potential regulatory functions (RegulomeDB score), and the Genomic Evolutionary Rate Profiling (GERP) score [55]. CADD scores of >10 are predicted to be the 10% most deleterious possible substitutions in the human genome, of >20 are predicted to be the 1%, and of >30 are predicted to be the 0·1%. RegulomeDB score is a categorical score from 1a to 7: SNPs showing the strongest evidence of being regulatory (affecting the binding of transcription factor) are given a score of 1, SNPs demonstrating the least evidence of being functional are given a score of 6, and 7 means that those SNPs do not have any annotations. GERP score ranges from −12·3 to 6·17, where higher scores indicate higher evolutionary constraint and a score greater than 2 can be considered constrained. To further investigate the functional role of genes most associated with RER values, we searched for known functions in the gene database of the National Center for Biotechnology Information (https://www.ncbi.nlm.nih.gov/gene/); each gene name, and its potential aliases (according to GeneCards, https://www.genecards.org/), was also entered into a PubMed search through the following algorithm: (GENE) AND ((malaria) OR (Plasmodium) OR (spleen) OR (red blood cell) OR (erythrocyte) OR (smooth muscle) OR (smooth muscular cell) OR (endothelium) OR (endothelial cells) OR (macrophage)) to identify any known association between these genes, malaria, and processes involved in RBC production or clearance, with a focus on cellular and biomechanical control of red blood cell deformability. In case of association between the considered gene and endothelial physiology, search was focused on articles mentioning the spleen. Regarding association with macrophage, we focused on articles mentioning their phagocytic function. We ranked genes according to the number and relevance of potential links between known gene functions and observed phenotypic features (namely, RBC deformability, splenomegaly, and innate processes involved in the quality control of RBC), as follows: no link (red boxes in Supplementary Table 4), potential and indirect link (orange boxes), direct link (green boxes). The relative abundance of each gene transcript in the spleen was estimated through RPKM, retrieved from the NCBI database.

### Gene-Based Association Testing

Gene-based association analysis was conducted using MAGMA through FUMA [50]. To identify candidate genes associated with RBC deformability (assessed by RER values adjusted for relevant covariates), the mean association of all SNPs within a gene was calculated, accounting for LD. Gene windows were extended 2kb upstream and 1kb downstream of the annotated gene start and end sites to include regulatory regions.

### Statistics

Binary variables were compared between groups using Chi-squared or Fisher’s exact test, as appropriate. Continuous variables were compared through Student’s t test or Mann-Whitney test, when comparing 2 groups; and through ANOVA or Kruskal-Wallis test, when comparing more than 2 groups, as appropriate. Multivariate analyses were performed using regression models, logistic or linear according to the dependent variable. Variables with p-value < 0·2 in univariate analysis were included in multivariate models, along with variables with potential strong interaction with the results, based on pathophysiological findings. All tests were 2-sided and a p-value lower than 0·05 was considered significant. Analyses were performed with GraphPad Prism 6©, R© (version 3.4.4), and RStudio© (version 1.1.442) softwares.

### Study approval

The BAObAB (Biocultural AdaptatiOn to malaria in Atacora, north Benin) research program was reviewed and approved by the ethics committee of the Research Institute of Applied Biomedical Sciences (CER-ISBA/Institut des Sciences Biomédicales Appliquées) in Benin (No 61/CER/ISBA/15). It received support from the Ministry of Health of Benin and has been carried out in close collaboration with the National Malaria Control Programme. All study participants were informed during meetings conducted in villages before the beginning of the study. Individual written informed consent was obtained from adults or from the parents or guardians of children, using a consent form translated in the subject’s native language (Ditamari, Bariba or Fulani).

## Results

### Splenomegaly is more frequent in Fulani

In a cross-sectional study, we examined 521 subjects (mean age, 21·9 +/- 17·7 years; range, 3 months-72 years; 48% female) of whom 157 (30·1%) were Fulani, 86 (16·5%) Gando, 159 (30·5%) Bariba, and 119 (22·8%) Otamari. For clarity, Gando, Bariba and Otamari were generally pooled in the non-Fulani group. Splenomegaly was more prevalent in Fulani than in non-Fulani (40·1% vs 23·5%; p=0·003; Figure 1a). There was a not statistically significant trend towards more prevalent anemia in Fulani (60·2% vs 51·5%, p=0·12; Figure 1b). Fever was rare in Fulani and non-Fulani (1·3% vs 2·5%; p=0·52). There was also a trend towards a more prevalent parasite carriage in Fulani (63·8% vs 56·6% for rapid diagnostic test (RDT) positivity p=0·14, and 44·9% vs 42·1% for thick smear p=0·57, Figure 1c-d). Median *P. falciparum* densities were insignificantly lower in Fulani than in non-Fulani (2500 vs 3440/μl, p=0·28, Figure 1 e1-3). Antimalarials had been used in the last 30 days by 7·04% and 11·23% of Fulani and non-Fulani subjects, respectively (p=0·17). In a multivariate analysis of factors influencing splenomegaly, we observed a strong association with the Fulani ethnicity (p=0·0007), the other significantly associated factors being young age, low hemoglobin and RDT positivity (Supplementary Table 1). On a longitudinal analysis of 6 cross-sectional studies performed in the same cohort from November 2015 through December 2017, the proportion of 113 subjects who remained RDT- negative across time was greater in Fulani than in non-Fulani.

**Figure 1.**
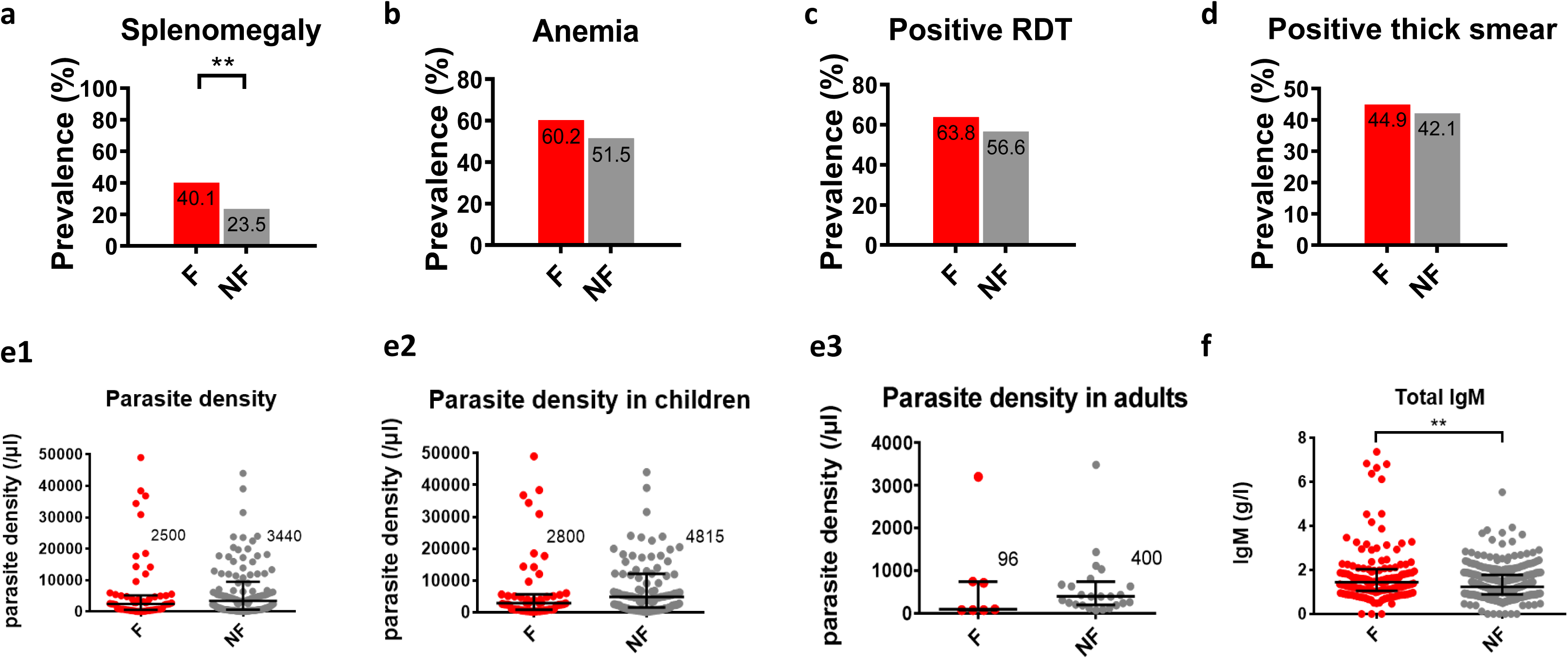
Splenomegaly is more prevalent in Fulani than in subjects from other ethnic groups. (a-d): Prevalence (%) of **(a)** splenomegaly, **(b)** anemia according to the WHO definition, **(c)** positive rapid diagnostic test for malaria and **(d)** positive thick smears in Fulani (F) and non-Fulani (NF). **(E1)** *Plasmodium falciparum* density (median, IQR) in Fulani and non-Fulani, shown separately in children **(e2)** and adults **(e3)** on thick smears (only positive values are shown). **(f)** Concentration of total IgM (median, IQR) in plasma from Fulani and non-Fulani. *: 0·01<p value<0·05; **: 0·001<p value<0·01. No symbol is shown if the difference is not significant. Categorical variables were compared through Chi-squared of Fisher’s exact test, as appropriate; parasite densities and total IgM values were compared through Mann-Whitney test.

### Circulating RBC are more deformable in Fulani than in other ethnic groups

Ektacytometry was performed on RBC from 248 randomly selected subjects (108 Fulani, 140 non- Fulani; 48·4% adults). Median elongation indexes of RBC measured at a shear stress of 30 Pascals (Figure 2a) were higher in Fulani than in non-Fulani (0·595 vs 0·589, p=0·039). Differences were also significant when comparing the four ethnic groups separately (p=0·0057; Supplementary Figure 1). Upon univariate analysis, splenomegaly and ethnicity were significantly associated with the Elongation Index of RBC (p=0·026 and p=0·013, respectively; Table 1). Upon multivariate analysis, only ethnicity remained significantly associated with the Elongation Index (p=0·016; table 1).

**Figure 2.**
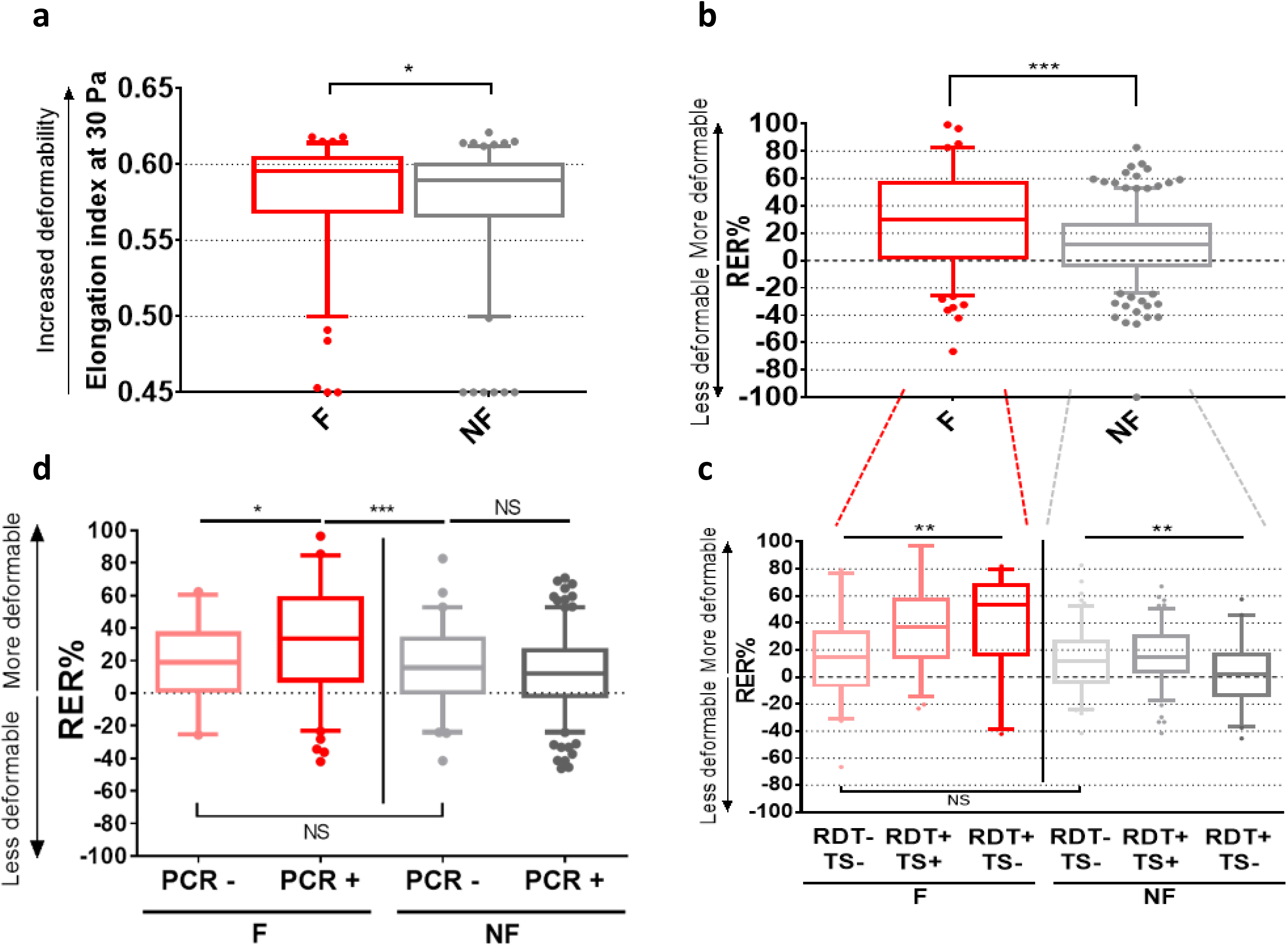
Circulating RBC are more deformable in Fulani than in subjects from other groups, and markers of recent *P. falciparum* infection are associated with enhanced RBC deformability in Fulani. (a) Elongation index (EI) of RBC measured by rotational ektacytometry at 30 Pa in Fulani (F) and non-Fulani (NF) (n=248). Higher EI values indicate enhanced RBC deformability. **(b)** Retention or enrichment rates (RER%) of circulating RBC from Fulani (F) and non-Fulani (NF) determined by filtration through microsphere layers (microsphiltration) (n=463). Positive RER (i.e., enrichment in test RBC downstream from filter) indicate enhanced RBC deformability (n=463). **(c)** RER determined by microsphiltration in Fulani (n=131) and non-Fulani (n=282) subjects with both RDT and thick smear negative for malaria parasites (RDT- TS-), or with both tests positive (RDT+ TS+) or with only RDT positive (RDT+ TS-). **(d)** RER values in Fulani (n=136) and non-Fulani (n=271) when considering subjects with negative or positive PCR for *P. falciparum*. Box plots indicate median and IQR values; bars indicate 5th and 95th percentiles. NS: no significant difference; *: 0·01<p value<0·05 ; **: 0·001<p value<0·01 ; ***: p value<0·001. Ektacytometry data were analyzed through Mann-Whitney test; Microsphiltration data were analyzed through Student’s t-test and ANOVA, as appropriate.

**Table 1.**
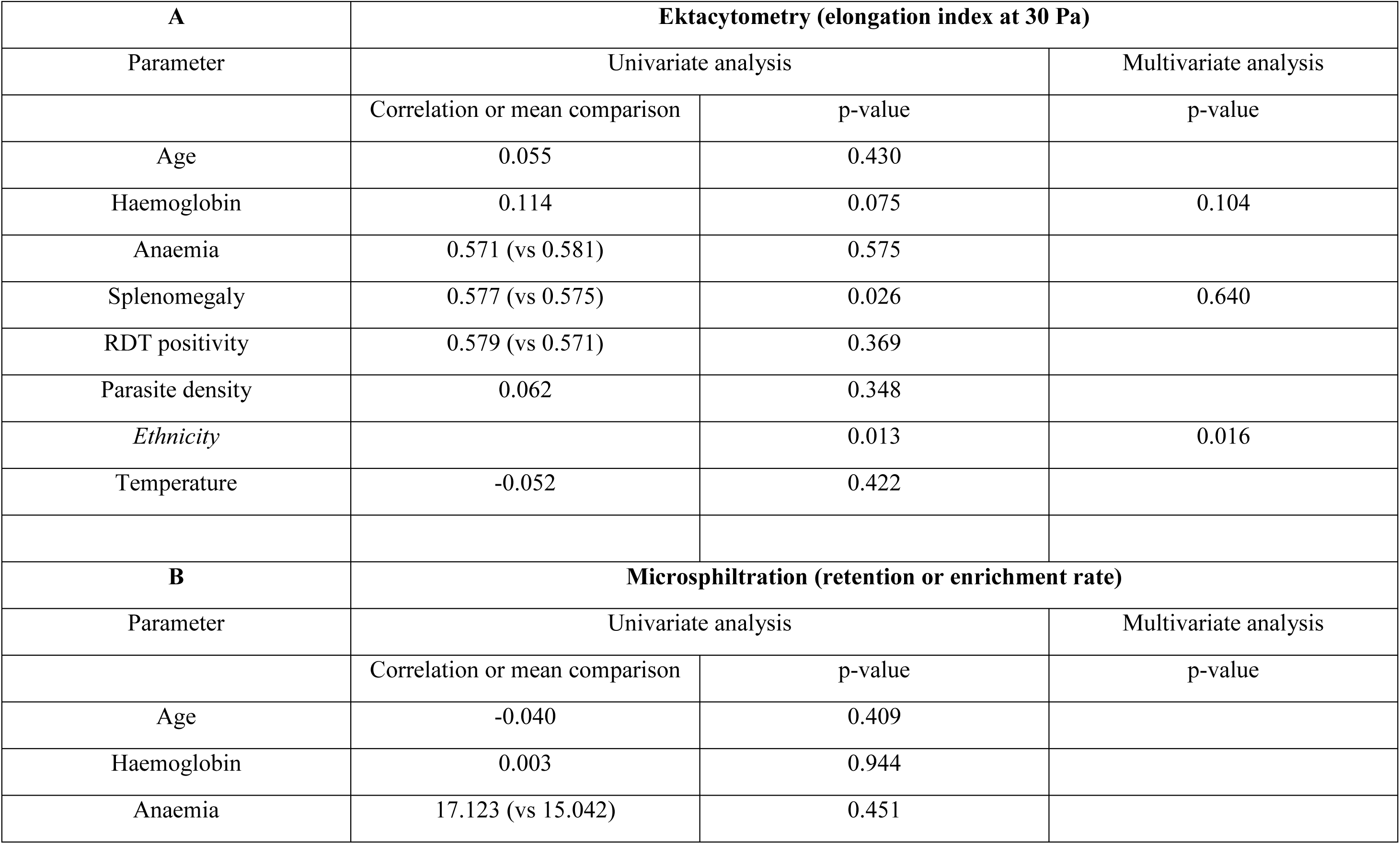
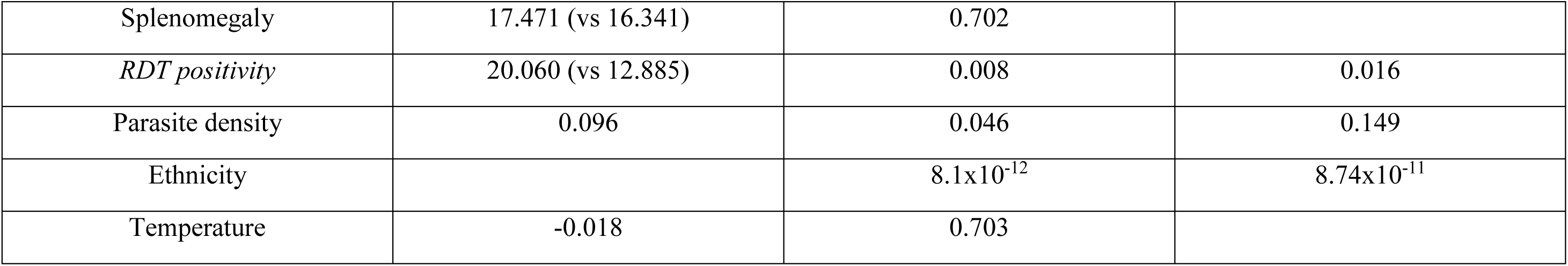
Univariate and multivariate analysis of factors associated with RBC deformability. assessed by ektacytometry (**A**: upper panel) or microsphiltration (**B**: lower panel). Variables with a p-value of 0.2 or less in univariate analysis were included in the multivariate analysis (multiple linear regression). Variables with a significant association with deformability measures after multivariate analysis are displayed in italic.

| **A** | **Ektacytometry (elongation index at 30 Pa)** |  |  |
| --- | --- | --- | --- |
| Parameter | Univariate analysis |  | Multivariate analysis |
|  | Correlation or mean comparison | p-value | p-value |
| Age | 0.055 | 0.430 |  |
| Haemoglobin | 0.114 | 0.075 | 0.104 |
| Anaemia | 0.571 (vs 0.581) | 0.575 |  |
| Splenomegaly | 0.577 (vs 0.575) | 0.026 | 0.640 |
| RDT positivity | 0.579 (vs 0.571) | 0.369 |  |
| Parasite density | 0.062 | 0.348 |  |
| *Ethnicity* |  | 0.013 | 0.016 |
| Temperature | -0.052 | 0.422 |  |
| **B** | **Microsphiltration (retention or enrichment rate)** |  |  |
| Parameter | Univariate analysis |  | Multivariate analysis |
|  | Correlation or mean comparison | p-value | p-value |
| Age | -0.040 | 0.409 |  |
| Haemoglobin | 0.003 | 0.944 |  |
| Anaemia | 17.123 (vs 15.042) | 0.451 |  |
| Splenomegaly | 17.471 (vs 16.341) | 0.702 |  |
| *RDT positivity* | 20.060 (vs 12.885) | 0.008 | 0.016 |
| Parasite density | 0.096 | 0.046 | 0.149 |
| Ethnicity | | $8.1 \times 10^{-12}$ | $8.74 \times 10^{-11}$ |
| Temperature | -0.018 | 0.703 |  |

Microsphiltration was performed on blood samples from 463 subjects (149 Fulani, 314 non-Fulani; 47·1% adults). For the 16 microplates used, negative control values were within the expected +/- 20% range, with a mean (IQR) of -7·66% (-13·91%; -6·71%). All positive controls displayed retention rates > 75% with a mean (IQR) value of 96·47% (95·87%; 99·66%). Retention or Enrichment Rates (RER) were significantly and markedly different between ethnic groups. Fulani had higher median RER, indicating enhanced deformability of their circulating RBC, compared to non-Fulani (30·1% vs 11·73%, p<0·0001; Figure 2b). RBC deformability was highest in Fulani, lowest in Bariba and intermediate in Gando and Otamari (Supplementary Figure 1). In univariate analysis, RER was influenced by ethnicity, parasite density and RDT positivity (p=8·1x10^-12^, 0·046 and 0·008, respectively); in the multivariate analysis, factors still significantly affecting deformability were ethnicity and RDT positivity (p=8·74x10^-11^ and 0·016, respectively) (Table 1). In summary, circulating RBC were more deformable in Fulani than in subjects from other ethnic groups. Two different methods, microsphiltration and ektacytometry, displayed consistent results. In the multivariate analysis, apart from ethnicity, the only factor associated with the deformability of circulating RBC was a positive RDT.

### Markers of infection with P. falciparum in Fulani are associated with a greater deformability of circulating RBC

Because differences were greater with microsphiltration than with ektacytometry we focused further analyses on results obtained with this method, which had also been performed on a larger number of samples. When considering only subjects with a negative RDT, RBC deformability was mildly (and insignificantly) higher in Fulani than in non-Fulani (median RER: 14·04% vs 11·72%, p=0·46). In subjects with a positive RDT, RBC deformability was markedly and significantly higher in Fulani than in other ethnic groups (median RER: 38·70% vs 11·96%, p<0·001). Within Fulani, RBC deformability was markedly greater in RDT-positive than in RDT-negative subjects (38·70% vs 14·04%, p=0·0007).

To analyse how markers of infection impact the deformability of circulating RBC, we then compared results from three groups, i.e. subjects in whom RDT and thick smears were both negative (a surrogate for absence of infection), subjects in whom RDT and thick smear were both positive (a surrogate for ongoing infection), and subjects in whom only the RDT was positive (a surrogate for recently resolved infection [56] or of major splenic tropism of parasitized RBC [23, 24]). The very few subjects (4 Fulani and 7 non-Fulani) in whom only thick smear was positive were not included in this analysis. In the context of this framework, marked differences were observed between Fulani and other ethnic groups (Figure 2c). In the Fulani group, RBC deformability was lowest in subjects with negative markers of infection, intermediate in subjects with markers both positive (thick smear and RDT), and highest in subjects with a positive RDT and a negative thick smear (median RER: 16·63%, 36·97% and 53·55% for both negative, both positive, and only RDT positive subjects, respectively, p=0·0022). By contrast, markers of infection correlated only mildly with RBC deformability in other ethnic groups (Figure 2c). PCR performed on a 407 subjects showed similarly that circulating RBC are more deformable in PCR-positive than in PCR-negative Fulani (RER=33·29% vs 18·91%, p=0·02, Figure 2d). No significant difference was observed when comparing PCR-negative and PCR-positive subjects from non-Fulani groups (RER=11·85% vs 15·55%, p=0·19, Figure 2d).

### Serum concentrations of total IgM are higher in Fulani than in non-Fulani

Total plasma IgM values (Figure 1f) were determined in 421 subjects. Median values were 1·41 g/L, and 1·57 g/L in non-Fulani and Fulani, respectively (p=0·009).

### A strong genetic component determines RBC deformability in Fulani

In a model incorporating household effects, the global heritability (h^2^) of RBC deformability for the whole sample of 420 subjects was 0·36 (p = 0·007). A stratified analysis by ethnicity revealed striking differences between Fulani and non-Fulani, with a very high and highly significant heritability estimate in the 135 Fulani (h^2^ = 0·71, p = 0·0028) and a near zero and nonsignificant one in the 285 non-Fulani (h^2^ =0·01, p=0·49) (Figure 3a). The variance component attributable to shared household environmental factors (c^2^) was only significant in non-Fulani (c^2^ = 0·30, p = 0·02). These results are consistent with the patterns of phenotypic correlations among relative pairs observed in Fulani, with correlations being highest between first-degree relatives, with similar correlation values between siblings and parent–offspring pairs (r=0·65, p < 10^-5^ and r=0·51, p = 0·005, respectively), lower but still significant between second-degree relatives (r= 0·37; p = 0·015) and even lower and not significant between more distant relatives (r=0·23; p=0·22) (Supplementary Table 2). This supports the existence of a strong, mainly additive, genetic component determining the deformability of circulating RBC in Fulani.

**Figure 3.**
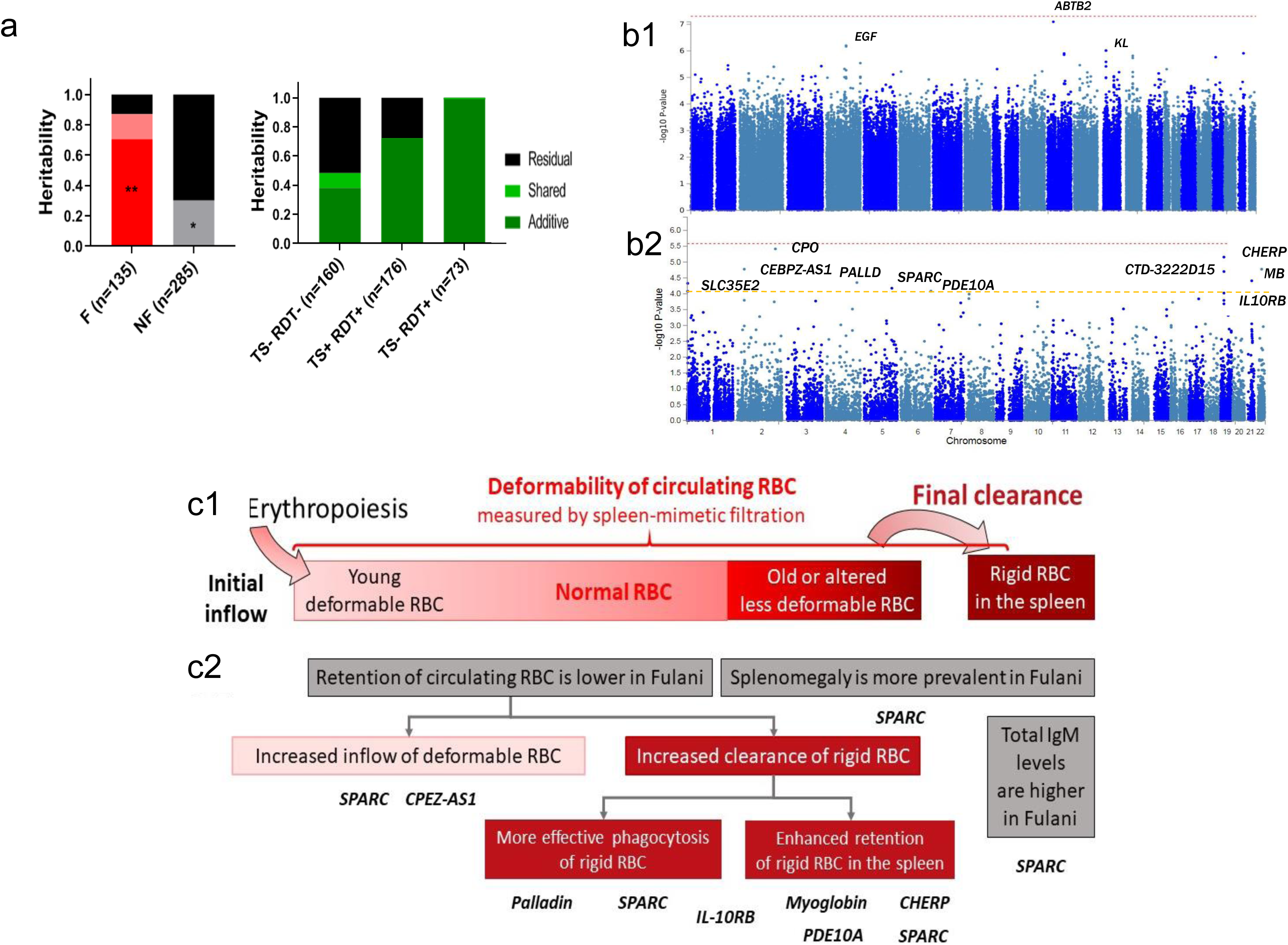
(a) Heritability of splenic RBC filtration. Using a variance component approach, the heritability of the deformability of circulating red blood cells (RBC) was determined either in relation with ethnicity in Fulani (F) or non-Fulani (NF) (left) or in the whole population depending on the status of *P. falciparum* infection (right). Additive heritability is estimated as the ratio of the additive genetic variance component to the total phenotypic variance. P-values were estimated using likelihood ratio tests. n: number of subjects; *: 0·01<p value<0·05 ; **: 0·001<p value<0·01 ; ***: p value<0·001; TS: thick smear; RDT: malaria rapid diagnostic test. **(b): Genome-wide association analysis for RBC deformability. (b1)** Each point of the Manhattan plot shows a single-marker variant, with genomic positions across chromosomes on the *X*-axis, and the association level corrected by genomic control on the *Y*-axis. The red dashed line shows the genome-wide significant *P*-value of 5 × 10^−8^. **(b2):** Gene-based test as computed by MAGMA. Each point of the Manhattan plot indicates a gene. Input SNPs were mapped to 19185 protein-coding genes. Genome wide significance (red dashed line in the plot) was defined at P = 0·05/19185 = 2·606 × 10^-6^. **(c1)** Determinants of retention upon splenic filtration of circulating RBC. **(c2)** Malaria-related phenotypic features observed in Fulani (grey boxes), potential mechanisms underlying the increased retention of circulating RBC in Fulani (red boxes), and genes potentially involved in these mechanisms as per literature analysis, the specific mechanisms being indicated in details in Supplementary Table 4 (Bold letters).

**Figure 4.**
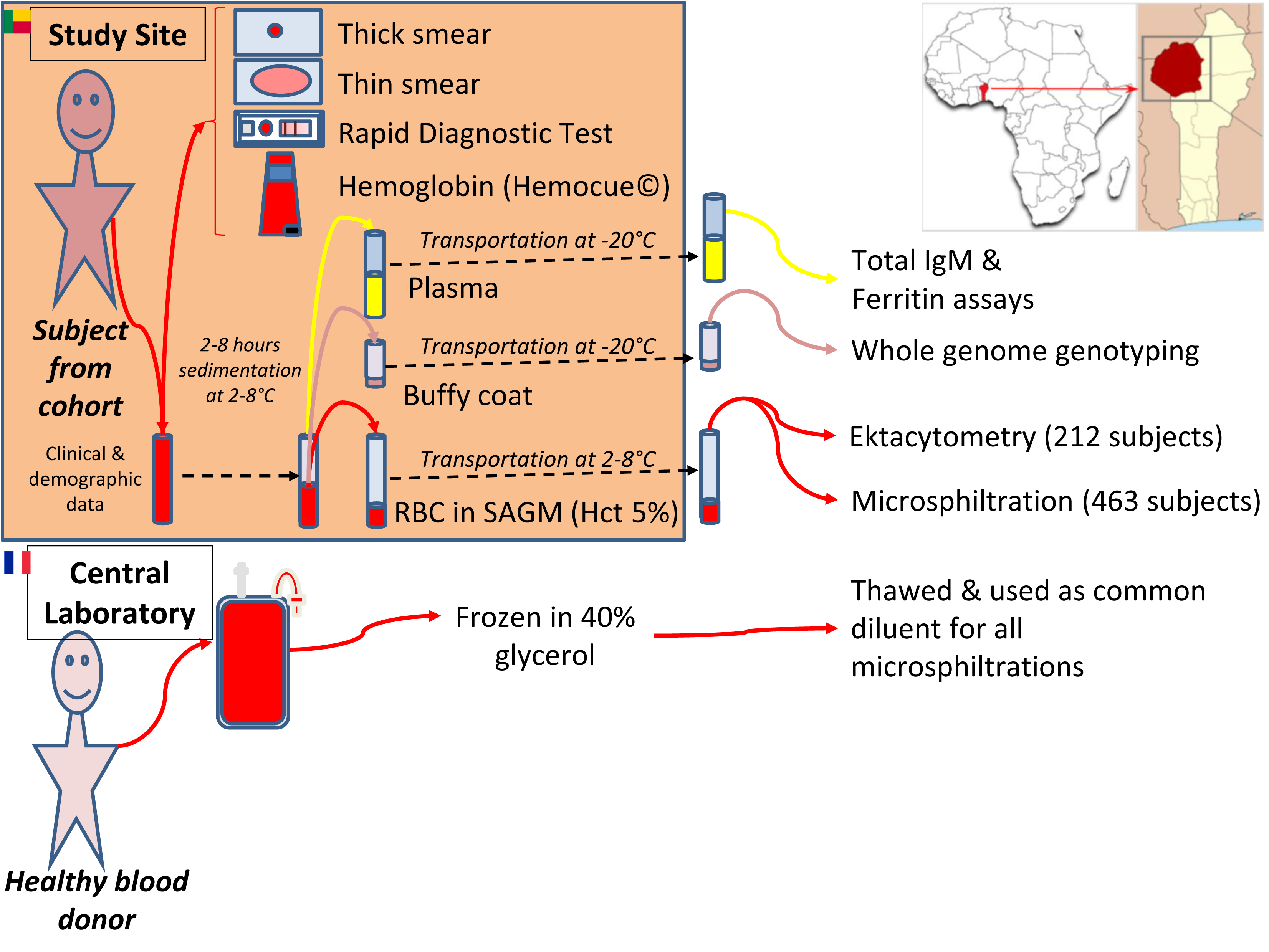
Overview of experimental procedures. Study location is indicated in the upper right corner (Birni district, Atacora department, Benin). SAGM: saline, adenine, glucose, mannitol. Ektacytometry was performed on 212 samples because of a shortage of polyvinypyrrolidone. PCR was performed on 424 samples because whole blood on filter paper could not be obtained in 97 subjects.

In the 160 subjects without markers of infection (thick smear and RDT both negative), the heritability of RBC deformability was almost identical to that observed in the whole cohort (h^2^ = 0·38, p=0·15) but raised to 0·72 (p=0·0002) in the 173 subjects with both markers positive, and to 0·99 with wide dispersion (p=0·07) in the 73 subjects with a positive RDT and a negative thick smear. This indicates that variations in the deformability of circulating RBC are under a strong influence of genetic determinants in Fulani and in subjects with positive markers of infection. The heritability of RBC deformability in Fulani with a positive RDT reached the maximum value of 1 (p<10^-5^) with a household effect estimated to be zero.

### Genome-wide association study of red blood cell deformability

Association of RBC deformability (through RER, adjusted for relevant covariates) with 2,281,899 high-quality SNPs, genotyped in 323 individuals, was evaluated genome-wide through a linear mixed model implemented in GenABEL [49]. The λ genomic inflation factor (ratio of the median of the observed distribution of the test statistic to the expected median) was 0·99997, consistent with a reliable adjustment for sample relatedness and population structure (Supplementary Figure 2). No variant reached the genome-wide significance threshold of 5×10^−8^ (Figure 3b1). However, three independent genomic loci on chromosomes 4, 11 and 13 showed suggestive association with RBC deformability (p-value ≤10^−6^), one of them being very close to the genome-wide significance threshold (SNP rs149514849 located at 11p13, close to gene *ABTB2*, p-value = 8·08 x 10^-8^; Table 2). None of the three lead SNPs in the three genomic risk loci were annotated as having predicted functional effects according to the three functional scores considered (CADD, RegulomeDB and GERP). However, according to ENSEMBL, the lead SNP rs149514849 is located on the binding site of a transcription factor, and has been associated with variations of the expression of CD44 in macrophages. CD44, a pleiotropic glycoprotein involved in cell-cell interactions, cell adhesion and migration, has a role in the phagocytic process of macrophages [57, 58]. Additionally, the lead SNP rs76559378 at 13q13 is in strong LD (*r*^2^ = 0·897) with the exonic *KL* SNP rs201936594 (same GWAS, *p* = 9·74 x 10^-7^) which displays a CADD score of 13·94, suggesting a likely deleterious effect. No other variant in the three associated genomic loci was annotated as functional. No significant association between our phenotype of interest and *ABTB2*, *EGF* and *KL* genes was found in the literature.

**Table 2.** Independent genomic risk loci suggestively associated (P-value ≤ 10-6) with RBC deformability (RER values adjusted for relevant covariates) in genome-wide association analysis. MAF, minor allele frequency; β is the estimate of the SNP effect under an additive model ; CADD, Combined Annotation-Dependent Depletion score; GERP, Genomic Evolutionary Rate Profiling.

| Genomic risk loci | Position (start-end in hg19) | No. GWAS SNPs | Lead SNP (rsID) | Non-effect allele | Effect allele | MAF | GWAS p-value | $\beta$ | SE | Location | Nearest gene | CADD score | RegulomeDB score | GERP score |
| --- | --- | --- | --- | --- | --- | --- | --- | --- | --- | --- | --- | --- | --- | --- |
| 11p13 | 34385832 | 1 | rs149514849 | C | T | 0.0211 | $8.08 \times 10^{-8}$ | -2.168 | 0.441 | Intergenic | *ABTB2* | 1.853 | 4 | -2.5 |
| 4q25 | 110854595-110907748 | 6 | rs528481745 | G | A | 0.0477 | $6.35 \times 10^{-7}$ | -1.653 | 0.362 | Intronic | *EGF* | 2.394 | 6 | -0.35 |
| 13q13 | 33567899-33590718 | 12 | rs76559378 | G | T | 0.0446 | $9.74 \times 10^{-7}$ | 1.560 | 0.347 | Promoter flanking region | *KL* | 0.729 | 7 | -2.34 |

To link the variants associated at p < 5 x 10^-6^ to genes, we applied three gene-mapping strategies implemented in FUMA [50]. These three methods identified a total of 25 genes (12, 7 and 11 by positional mapping, eQTL mapping and chromatin interaction mapping, respectively), three of which were identified by two mapping methods (*CHERP*, *CTD-3222D19* and *ABCC1*) and only one (*SLC35E1*) by the three (Supplementary Table 3). A thorough literature review did not find any relevant association between any of these 4 genes and mechanisms potentially underlying the newly defined phenotype. Regarding the remaining 21 genes, few associations were found in the available literature, all for genes identified through chromatin interaction mapping: *CD59* expression is decreased on *Plasmodium*-infected erythrocytes, which may promote erythrophagocytosis [59]; *LMO2* plays an important role in erythropoiesis [60]; and *IL1RAP* upregulates the expression of the “don’t eat me” signal CD47 to inhibit macrophage phagocytosis [61].

Gene-based association testing was then performed with MAGMA [62], which uses the mean association signal from all SNPs within each gene while accounting for LD. A total of 19,185 genes were analyzed. A multiple testing-corrected *p* value threshold of 2·61 × 10^−6^ was applied to identify genes significantly associated with RBC deformability. No gene score was above this significance threshold (Figure 3b2; see the corresponding quantile–quantile plot in supplementary figure 2b). Nevertheless, we analysed the top ten genes that displayed p-values between 3·84x10^-6^ and 8·25x10^-5^ (Figure 3b2 and Supplementary Table 4). A thorough search and analysis of available information did not identify any link between the selected genes and malaria, excepted for *IL10RB*, which has been associated with severe malaria in one study [63]. Two genes, *PALLD* [64] and *SPARC* [65] have a role in erythropoiesis, that may indirectly affect the deformability of circulating RBC. Regarding macrophage-related processes and innate immunity, PALLD is a regulator of phagocytosis [66] and *SPARC* has a role in development of autoimmunity, especially B cell-mediated [67], *IL10RB* is directly associated with the human immune response. Not least, five genes, *CHERP*, *MB*, *PALLD*, *PDE10A*, and *SPARC* are associated with components of the splenic red pulp where RBC clearance occurs: *CHERP* may be involved in the regulation of excitation-contraction coupling through an interaction with RyR1 [68]; myoglobin regulates NO production in vascular smooth muscle [69], *PALLD* is involved in mesenchymal and muscle cells formation, maturation, migration, and contraction [70], *PDE10A* may have a role in the regulation of cAMP and cGMP and hence contraction of smooth muscle [71, 72], and *SPARC* is expressed in mice by cell forming the wall of red pulp venules where the filtration of RBC takes place [73]. We considered genes involved in contraction because the contraction of smooth muscle cells and myofibroblasts potentially regulates slits size and RBC retention upstream from sinus walls in the spleen red pulp [74]. According to the Gene database of the National Center for Biotechnology Information, 5 of the top 10 genes (*CHERP, CEBPZ-AS1, IL10RB, PALLD, SPARC*) displayed a significant splenic expression (RPKM values ranging from 10·66 to 123·8). Immunohistochemistry performed on a sample of human spleen showed the presence of SPARC within the red pulp, especially at the sinus wall (Supplementary Methods and Supplementary Figure 3) and in smooth muscular cells of arteries.

## Discussion

Malaria is an archetypical example of how an infectious disease has shaped the human genome, and hemoglobin S mutation is the first genetic trait robustly shown to innately protect against a microbial agent [1]. Natural resistance to malaria however may extend well beyond RBC polymorphisms and we show here that a peculiar malaria-related phenotype observed in African subgroups likely results from spleen-related mechanisms. Splenomegaly, anemia and hyper-IgM are indeed prevalent in Fulani and we found their circulating RBC to be more deformable than in other groups. These original results were obtained by filtering RBC in a spleen-mimetic way and no known RBC-related trait was associated with this new cell-based, highly inheritable phenotypic trait. Our preliminary genome-wide association study points to several genes expressed in the spleen and potentially involved in the clearance of RBC, one of which, *SPARC*, has previously been linked to splenomegaly and IgM production. Taken together, these results suggest that the mechanisms underlying the peculiar, “malaria-hyperreactive” phenotype of Fulani, shared by other groups worldwide [75], may stem from innate processes driving more altered and infected RBC to the spleen, thereby increasing the antigenic stimulus to this immunologically reactive organ. Recent major observations [23, 24] in Indonesia have indeed shown that in asymptomatic carriers of malaria parasites, far more intact infected RBC are in the spleen than in the peripheral blood. The original reactivity to malaria in some subgroups may therefore be only partially immunological in essence, and also depend on microvascular and interstitial processes that regulate the filtration of RBC. Hyper-stringent filtration of RBC by the human spleen parsimoniously explains the original splenomegaly-anemia-Hyper-IgM syndrome of Fulani.

Assuming that mature RBC cannot autonomously modify their deformability, an increase in the deformability of circulating RBC can result either from an inflow of more deformable cells or from the clearance of the less deformable cells (Figure 3c1-2). When comparing parasite-positive with parasite-negative Fulani, the difference in the proportion of circulating rigid RBC was greater than 20%. In subjects with acute malaria attacks, the mean peak reticulocytes concentration is generally markedly lower than 7% [76–78]. Therefore, the inflow of new RBC is unlikely to explain the malaria-related enhancement of RBC deformability in Fulani who had in most case low-grade parasitemia. Acute malaria induces an inflow of infected and modified uninfected RBC in the circulation [22, 26] which are less deformable than normal RBC [25–27,29,31,79,80]. The measured hyperdeformability of circulating RBC observed in Fulani thus likely increases essentially in reaction to an enhanced clearance of the least deformable subset of these circulating RBC. The major organ that controls RBC deformability is the spleen. Several RBC labelling studies in humans [77,81–83] have shown that the clearance of RBC with either mechanical or surface alterations is enhanced in *P. falciparum*–infected subjects, very strongly so when splenomegaly is present [81]. The “hungry” post-malarial spleen even displays an increased clearance of normal RBC [78, 83]. Taken together, these observations suggest that, in Fulani, splenic filtration of RBC is enhanced, with strong association with the presence of parasite antigens in circulation. Interestingly, when malaria parasites intensely accumulate in the spleen, as recently uncovered [23, 24], very few parasites circulates but parasite antigens are released in plasma and captured by RDT.

If Fulani display indeed a particularly stringent splenic filtration of RBC, this not only explains the higher deformability of their circulating RBC but also partial antimalarial protection, and the more prevalent anemia and splenomegaly, three hallmarks of the malaria-related ‘Fulani phenotype’ [5]. This new ‘stringent splenic filtration’ phenotypic trait is expected to be partially protective by clearing part of the circulating parasite load [29] and pathogenic by precipitating splenomegaly and anemia [22, 30]. The deformability of ring-infected and that of uninfected RBC overlap [25], so that protective retention of a high proportion of infected RBC would also precipitate anemia, caused by an enhanced retention of uninfected RBC in the spleen. Enhanced retention of RBC also cause congestive splenomegaly observed indeed in a recent splenectomy study in a malaria-endemic area (Kho *et al.*, in preparation). This accumulation of infected RBC in the spleen of Fulani perfectly matches the frequent discrepancy between a positive RDT and negative thick smear displayed by subjects who had the most deformable RBC in circulation, reflecting the most stringent splenic retention. Splenic parasites release indeed antigens visualized by RDT but do not circulate hence the negative smear. Not least, by dispatching more parasites to the responsive splenic environment [84], malaria-enhanced splenic filtration of infected RBC would also explain the strong immunological stimulation observed in Fulani [4,6,9,18,19]. Higher circulating IgM levels observed previously [8] and confirmed in this study, fit a predominantly splenic stimulation, as many IgM-producing cells are in the spleen [85]. Interestingly, antibody responses against *Helicobacter pylori* or rubella virus are not greater in Fulani than in other groups [8] suggesting that the enhanced stimulation in this group is specific to *Plasmodium.* The mechanism of this microbe-specific response had not been identified yet and is explained by our model. Along these lines, the proposed ‘stringent splenic filtration’ trait is reminiscent of hyper- reactive malarial splenomegaly [3], a syndrome defined by gross splenomegaly, elevated antibody concentrations to *P. falciparum*, low parasitemia, and elevated total IgM levels [86]. Hyper- reactive malarial splenomegaly is more frequent in Fulani than in other groups [3,5,7], and may be the chronic complication of repeated episodes of hyper-reactive splenic filtration in infected subjects. Chronic antigenic stimulation of the spleen is a convincing explanation for the link between hyper-reactive splenomegaly and marginal zone splenic lymphoma [87]. Our phenotypic observations in Fulani are however milder than HMS, as only very few subjects had a full-blown HMS with gross splenomegaly and very high IgM levels.

This study is not devoid of weaknesses and some of our assumptions would benefit from more direct demonstrations. We provide no direct experimental evidence that the malaria-enhanced clearance of less deformable RBC is operated by the spleen. We nevertheless favour a spleen related process because ektacytometry results were significant only at 30 Pa, a surrogate for shear stress in the splenic red pulp [36], and because ethnicity- and infection-related differences in RBC deformability were the most pronounced with microsphiltration, a mimic of splenic retention of RBC [31]. This assumption is also based on reports of experimental transfusion of malaria patients with labelled RBC showing increased RBC retention during or after malaria attacks [81–83]. Not least, that several genes selected by our preliminary GWAS approach are expressed in the spleen, which strengthens the suspicion of a spleen-related process. The mechanisms underlying the putative malaria-enhanced splenic hyperfiltration in Fulani are currently unknown and difficult to explore directly. Contrast-enhanced ultrasonography can quantify the proportion of splenic blood flow to the slow, filtering compartment [29] but can hardly be performed in asymptomatic naturally infected subjects in remote areas. Currently available mouse models of malaria are not adapted to these explorations. Mechanical processes affecting *Plasmodium*-infected RBC display marked differences between human and rodent parasite species, and humanized mice infected with *P. falciparum* have a defective splenic filtration, as illustrated by the presence of stiff mature asexual stages and immature gametocytes in circulation [88, 89]. Regulation of the width of inter- endothelial slits likely exists in humans [74, 90] but exploring it directly is beyond the scope of current experimental methods. Recently, natural selection on genetic variants in the *BDKRB2* and *PDE10A* genes has been shown to affect the human ability to withstand diving-related hypoxia likely through pooling of RBC in the spleen [91]. This strongly supports the existence of genetically-driven mechanisms that regulate splenic filtration of RBC. Of interest, *PDE10A* is among the top-10 genes revealed by gene-based testing in our preliminary GWAS.

Due to recruitment constraints (for safety reasons, North Benin has become incompatible with field studies since 2018), our GWAS did not reach a fully accurate power. Nevertheless, several genes warranting further investigations were identified. In keeping with previous reports in Fulani [92], none of the usual genetic polymorphisms associated with protection from malaria, such as hemoglobinopathies or blood groups, had high scores. Of the 10 genes the most significantly associated with RBC deformability in the gene-based tests, 5 (*CHERP*, *CEBPZ-AS1*, *IL10RB*, *PALLD*, *SPARC*) are highly transcribed in the spleen. Four selected genes (*CHERP*, *myoglobin*, *PALLD*, *PDE10A*) play a role in smooth muscular contraction and thus may regulate the dispatching of splenic blood flow to the slow filtering circulation of the spleen red pulp, as well as the opening of splenic inter-endothelial slits where altered and parasitized RBC are mechanically trapped [29,74,93]. SPARC has been involved in vascular smooth muscle cells proliferation [94], is expressed on splenic sinus cells (Supplementary Figure 3 and [73]) and impacts the production of collagen IV [95], a major component of filaments surrounding splenic sinuses [96]. Like other selected genes, it may therefore impact the regulation of RBC filtration by the spleen, but SPARC is also linked to the recruitment of macrophages, that eliminate altered RBC, and SPARC-null mice have an altered splenic marginal zone and B-cell response [73], including abnormal levels of IgM.

Hyper-reactive filtration of RBC by the spleen is the most parsimonious explanation for a new phenotypic trait in Fulani where, in addition to well-known over-prevalence of splenomegaly and anemia, we reveal that circulating RBC are more deformable, almost exclusively in subjects with markers of ongoing low-grade (likely intrasplenic) accumulation of parasites [23, 24], or recent parasite carriage. Beyond the understanding of the malaria-related phenotype in Fulani, infection- related splenic hyper-filtration of RBC may contribute to the pathogenesis of other RBC diseases. For example, splenic sequestration crises in sickle cell disease - the acute trapping of RBC in the spleen - often follow acute episodes of fever or infections [97]. The potential impact of RBC filtration on antigen dispatching in the spleen also deserves consideration as it may display a major impact on the understanding of immunization by intravascular pathogens.

## Author contributions

AS, AG, PB and BH designed the study; BH, GV, CR, P-AN, AF, CC, JS, C-HC, FA, EC, RA, NF, JC, AS and PB performed the experiments; HA, AS, AG, MG, FT, DC, JC, PB, and BH collected clinical data; FP, LG and AS analyzed genetic data; FS, MM and OH provided scientific insights to experimental design; BH and PB wrote the initial draft of the manuscript; all authors critically reviewed and approved the final version of the manuscript.

## Acknowledgements

We thank all the subjects who participated in the study; Michaël Dussiot, Mickaël Marin, Sylvestre Biligui and Nathalie Chartrel for outstanding technical support; our research team in Benin for excellent technical and logistical achievements: Séverin Vignigbe, Pépin Kounou, Arnaud Fara Sarre, Guerelle Gnamou, Sabi N’Dah Kouagou, Wilson Houssou, Francis Homegnon; Jehane Fadlallah for her help in the recruitment of control patients; Nicolas Bréchot for his logistical help; Geneviève Milon, Adrian Luty and Jacques Lebras for giving scientific guidance and participating in discussions during the course of this research. This research was supported by a grant from Labex GR-Ex and from the National Geographic Society Committee for Research and Exploration (Grant 9804-15). The labex GR-Ex, reference ANR-11-LABX-0051 is funded by the program Investissements d’avenir of the French National Research Agency, reference ANR-11-IDEX- 0005-02. It also received funding from IMEA (Institut de Médecine et d’Epidémiologie Appliquée) and IRD (Institut de Recherche pour le Développement); BH was supported by a grant from Région Ile de France (DIM Malinf ; grant dim150030).

## Data sharing statement

Deidentified data used in this study will be made available upon reasonable request and approval by Pierre A Buffet and Audrey Sabbagh. Requests can be made by email to the corresponding author.

