## Supplementary Figures for "Splenic clearance of rigid erythrocytes as an inherited mechanism for splenomegaly and natural resistance to malaria"

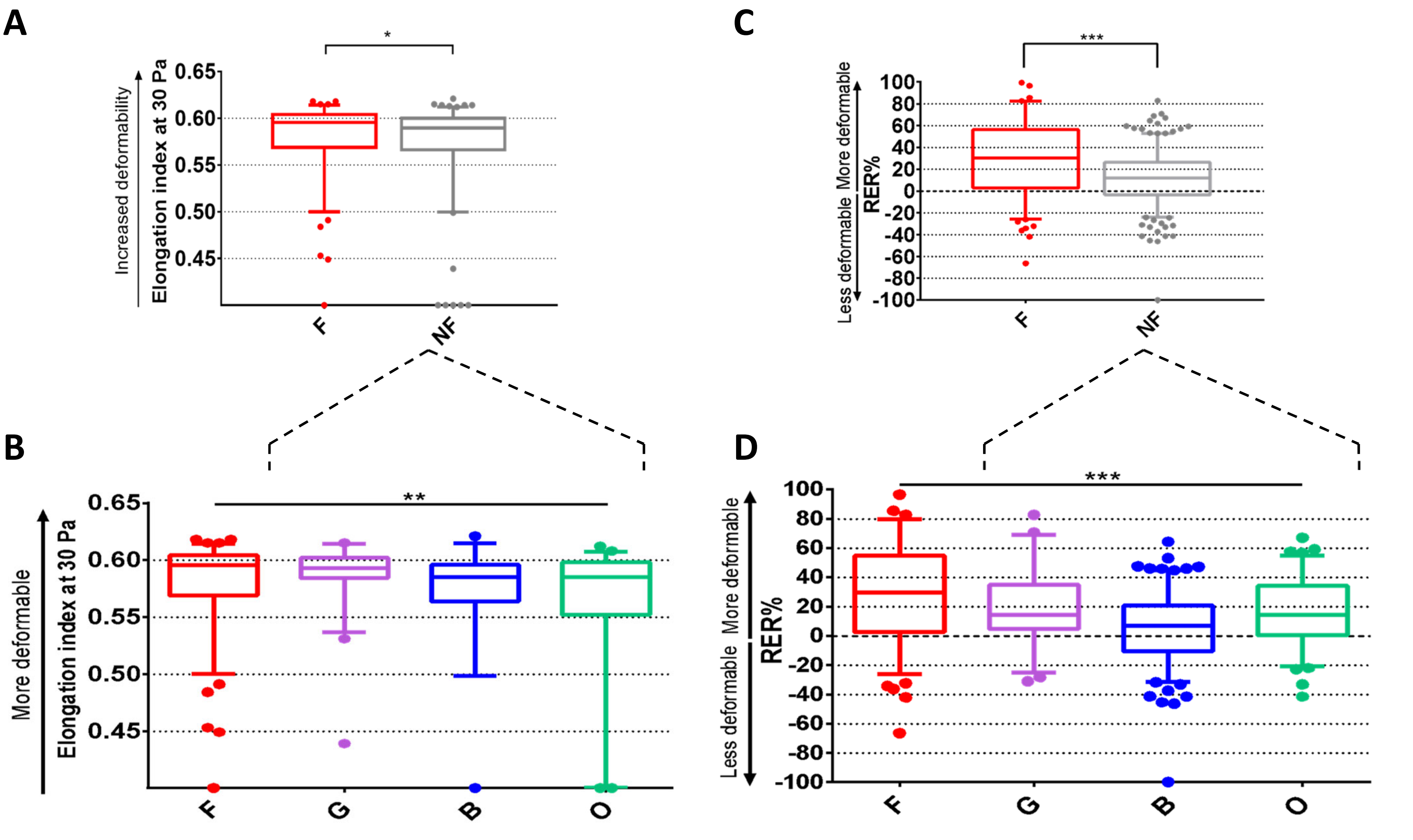


**Supplemental figure 1. Circulating RBC are more deformable in Fulani than in subjects from other groups**. **(A)** Elongation index (EI) of RBC measured by rotational ektacytometry at 30 Pa in Fulani and non-Fulani. Higher EI values indicate enhanced RBC deformability. **(B)** EI at 30 Pa of RBC from the different ethnic groups. **(C)** Retention or enrichment rates (RER%) of circulating RBC from Fulanis (F) and non-Fulanis (NF) determined by filtration through microsphere layers (microsphiltration). Positive RER (i.e., enrichment in test RBC downstream from filter) indicate enhanced RBC deformability. **(D)** RER in the different non-Fulani ethnic groups. G: Gando; B: Bariba; O: Otamari.. Boxes indicate median and IQR values. Bars indicate 5th and 95th percentiles. *: 0.01<p value<0.05; **: 0.001<p value<0.01; ***:p value<0.001. Microsphiltration data were analyzed through Student's t test and ANOVA, as appropriate; ektacytometry data were analyzed through Mann-Whitney and Kruskal-Wallis tests, as appropriate.

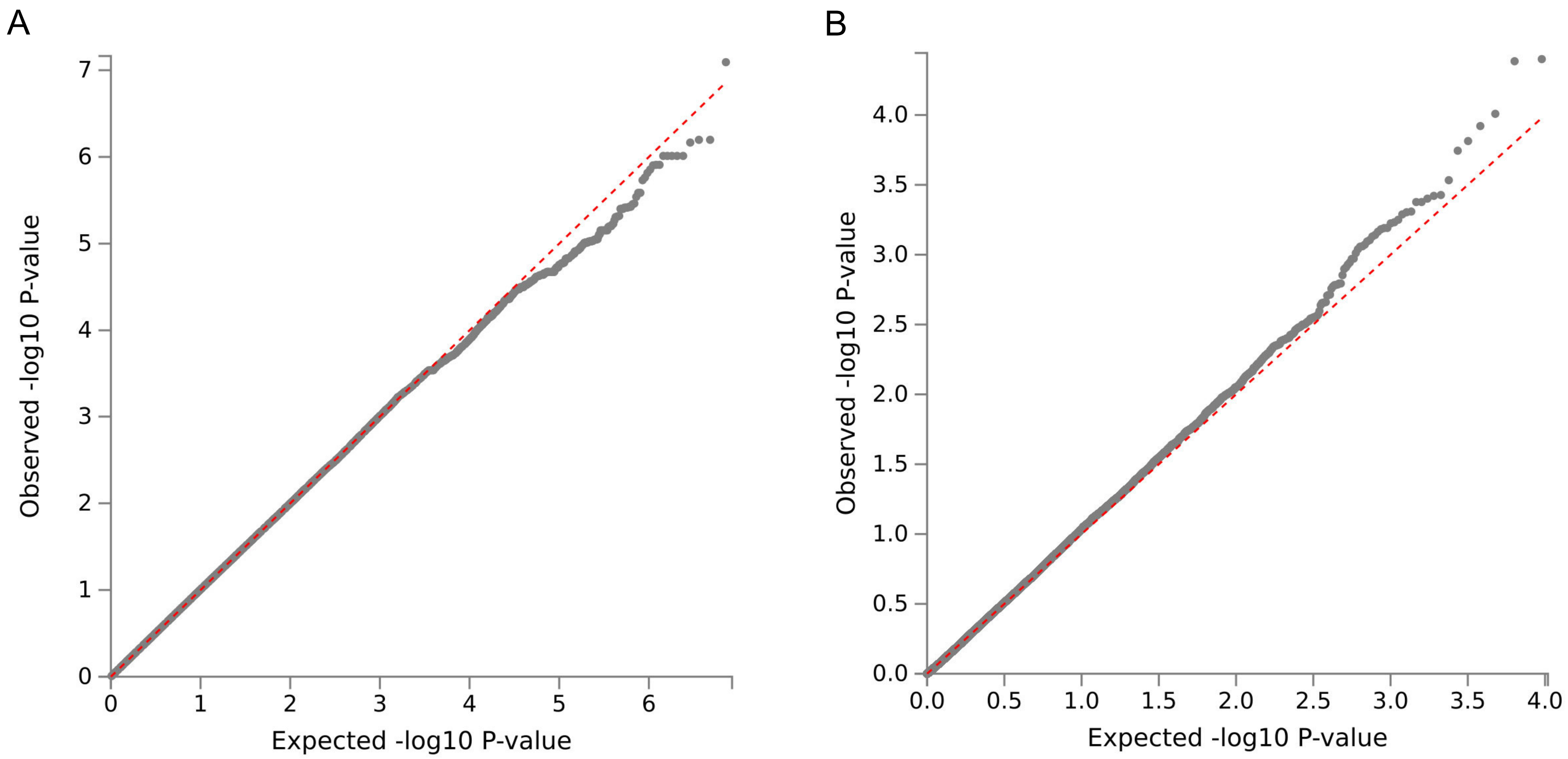


**Supplemental Figure 2**: **Quantile-quantile plots of *P*-values for genome-wide association analysis with RBC deformability (RER values adjusted for relevant covariates).** The red dashed line is the slope expected under no inflation and no true associations, the y-axis represents the observed $-\log_{10}$ of *P*-values, and the x-axis represents the expected $-\log_{10}$ of *P* values under the null hypothesis of no association. **(A)** Single-marker genome-wide association tests. **(B)** Gene-based tests as computed by MAGMA from single-marker association tests.

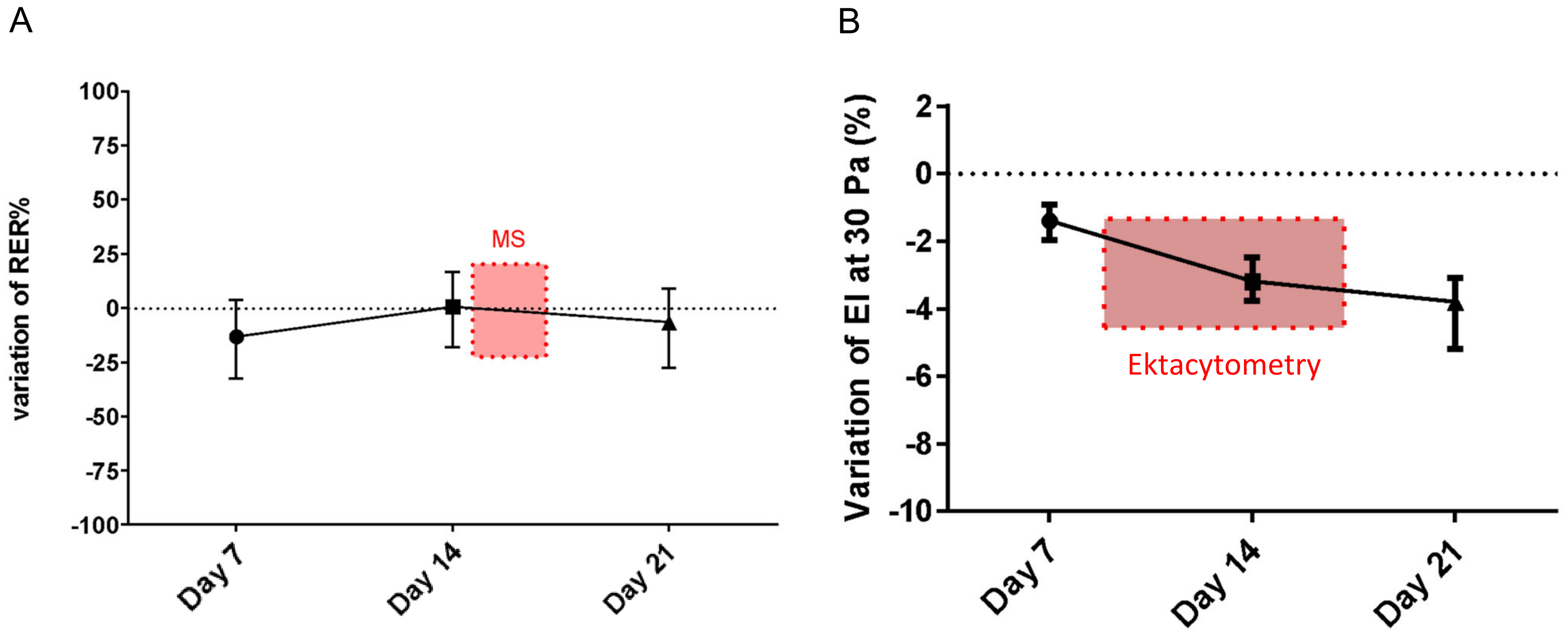


**Supplemental Figure 3**: **preservation of RBC in SAGM at 5% haematocrit allows delayed assessment of RBC deformability through microsphiltration and ektacytometry**. **(A)** Displayed are median and interquartile ranges of RER variation between day 0 and days 7, 14 and 21 post sampling in 18 healthy RBC donors. The mean RER variations were -0.9% at day 14 and -8.4% at day 21. The red rectangle indicates the timing of MS after collection of RBC in Benin. **(B)** In the same population, displayed are median and interquartile ranges of Elongation Index at 30 Pa variation between day 0 and days 7, 14 and 21 post sampling. The red rectangle indicates the timing of ektacytometry after RBC collection in Benin. MS : microsphiltration.
