## Supplementary Tables for "Splenic clearance of rigid erythrocytes as an inherited mechanism for splenomegaly and natural resistance to malaria"

| Variable | Test | Univariate | | Multivariate |
|---|---|---|---|---|
| | | Mean or proportion comparison (with vs without splenomegaly) | p-value | p-value |
| Ethnicity (proportion of Fulani, %) | $\chi 2$ | 25.73 vs 43.07 | 0.0003 | 0.0002 |
| Age (years) | Wilcoxon | 26.57 vs 7.89 | $< 2.2 \times 10^{-16}$ | $3.18 \times 10^{-6}$ |
| Haemoglobin (g/dl) | t test | 11.95 vs 9.89 | $< 2.2 \times 10^{-16}$ | 0.0005 |
| RDT positivity (%) | $\chi 2$ | 47.95 vs 81.02 | $1.42 \times 10^{-10}$ | 0.0615 |
| Temperature (°C) | t test | 37.02 vs 37.29 | $7.48 \times 10^{-9}$ | 0.019 |

**Supplemental Table 1. Univariate and multivariate (logistic regression) analysis of the clinical and biological parameters associated, in the whole studied population, with palpable splenomegaly in December, 2017**. RDT: rapid diagnostic test for malaria. Anaemia and Fever have been omitted because of redundancy with haemoglobin and temperature values.

| | Number of pairs (N) | r | +/- SE | p-value |
|---|---|---|---|---|
| Total population (422 individuals) | | | | |
| Parent :offspring | 337 | 0.47 | 0.06 | *** |
| Sibling | 138 | 0.27 | 0.12 | * |
| Total 1st degree | 475 | 0.41 | 0.04 | *** |
| Total 2nd degree | 122 | 0.24 | 0.13 | * |
| Total ≥ 3rd degree | 76 | 0.16 | 0.16 | |
| Fulani (135 individuals) | | | | |
| Parent :offspring | 106 | 0.65 | 0.08 | *** |
| Sibling | 55 | 0.51 | 0.15 | ** |
| Total 1st degree | 161 | 0.61 | 0.06 | *** |
| Total 2nd degree | 70 | 0.37 | 0.19 | * |
| Total ≥ 3rd degree | 66 | 0.23 | 0.20 | |
| Non-Fulani (285 individuals) | | | | |
| Parent :offspring | 231 | 0.29 | 0.08 | *** |
| Sibling | 83 | 0.18 | 0.11 | |
| Total 1st degree | 314 | 0.26 | 0.06 | *** |
| Total 2nd degree | 49 | 0.11 | 0.23 | |
| Total ≥ 3rd degree | 10 | 0.16 | 0.57 | |

**Supplemental Table 2. Adjusted intrafamilial correlation coefficients for the RBC deformability trait assessed by microsphiltration**. All correlation coefficients were adjusted for RDT positivity and ethnicity. r: intrafamilial correlation coefficient; SE: standard error; * $p < 0.05$; ** $p < 0.01$; *** $p < 0.001$.

| | | | | | Positional mapping | | | eQTL mapping | | | CI mapping |
|---|---|---|---|---|---|---|---|---|---|---|---|
| Gene | Position (chromosome: start-end) | Genomic risk locus | Min GWAS p-value | Lead SNP (rsID) | nSNPs | Max CADD score | Min RegulomeDB score | nSNPs | Direction | p-value | Tissue name |
| *CD59* | 11 : 33719807-33757991 | 11p13 | 8,08e-08 | rs149514849 | 0 | NA | NA | 0 | NA | NA | Mesenchymal Stem Cell |
| *LMO2* | 11 : 33880122-33913836 | 11p13 | 8,08e-08 | rs149514849 | 0 | NA | NA | 0 | NA | NA | Mesenchymal Stem Cell |
| *NAT10* | 11 : 34127149-34169217 | 11p13 | 8,08e-08 | rs149514849 | 0 | NA | NA | 0 | NA | NA | Mesenchymal Stem Cell |
| *ABTB2* | 11 : 34172535-34379555 | 11p13 | 8,08e-08 | rs149514849 | 1 | 1.853 | 4 | 0 | NA | NA | NA |
| *AC132216.1* | 11 : 33902189-33903338 | 11p13 | 8,08e-08 | rs149514849 | 0 | NA | NA | 0 | NA | NA | Mesenchymal Stem Cell |
| *EGF* | 4 : 110834040-110933422 | 4q25 | 6,35e-07 | rs528481745 | 6 | 4.387 | 6 | 0 | NA | NA | NA |
| *KL* | 13 : 33590207-33640282 | 13q13.1 | 9,74e-07 | rs76559378 | 5 | 13.94 | 4 | 0 | NA | NA | NA |
| *TTC3* | 21 : 38445526-38575413 | 21q22.13 | 1,23e-06 | rs148698956 | 2 | 8.98 | 4 | 0 | NA | NA | NA |
| *CHERP* | 19 : 16628700- | 19p13.11 | 1,73e-06 | rs113818017 | 15 | 5.9 | 4 | 1 | - | 2.81227e-12 | NA |

| | | | | | | | | | | | |
|---|---|---|---|---|---|---|---|---|---|---|---|
| | 16653341 | | | | | | | | | | |
| *SLC35E1* | 19 : 16660642-16683193 | 19p13.11 | 1,73e-06 | rs113818017 | 4 | 3.069 | 6 | 1 | + | 1.6696e-38 | Mesenchymal Stem Cell |
| *CTD-3222D19.2* | 19 : 16589878-16739015 | 19p13.11 | 1,73e-06 | rs113818017 | 24 | 5.947 | 5 | 0 | NA | NA | Mesenchymal Stem Cell |
| *SMIM7* | 19 : 16741562-16771253 | 19p13.11 | 1,73e-06 | rs113818017 | 0 | NA | NA | 17 | - | 3.1641e-07 | NA |
| *PLXNA2* | 1 : 208195587-208417665 | 1q32.2 | 3,52e-06 | rs1152834 | 3 | 4.431 | 5 | 0 | NA | NA | NA |
| *LEPREL1* | 3 : 189674517-189840226 | 3q28 | 3,76e-06 | rs710554 | 6 | 10.13 | 3a | 0 | NA | NA | NA |
| *CLDN16* | 3 : 190040330-190129932 | 3q28 | 3,76e-06 | rs710554 | 0 | NA | NA | 0 | NA | NA | Aorta; Mesenchymal Stem Cell |
| *MASP1* | 3 : 186935942-187009810 | 3q28 | 3,76e-06 | rs710554 | 0 | NA | NA | 0 | NA | NA | Mesenchymal Stem Cell |
| *CLDN1* | 3 : 190023490-190040264 | 3q28 | 3,76e-06 | rs710554 | 0 | NA | NA | 0 | NA | NA | Aorta; Mesenchymal Stem Cell |
| *IL1RAP* | 3 : 190231840-190375843 | 3q28 | 3,76e-06 | rs710554 | 0 | NA | NA | 0 | NA | NA | Mesenchymal Stem Cell |
| *PMS1* | 2 : 190649107-190742355 | 2q32.2 | 3,97e-06 | rs785259 | 0 | NA | NA | 7 | - | 3.7381e-08 | NA |
| *ASNSD1* | 2 : 190526111-190535557 | 2q32.2 | 3,97e-06 | rs785259 | 0 | NA | NA | 7 | + | 8.7163e-32 | NA |
| *C2orf88* | 2 : 190744335-191068210 | 2q32.2 | 3,97e-06 | rs785259 | 8 | 8.998 | 6 | 0 | NA | NA | NA |

| | | | | | | | | | | | |
|---|---|---|---|---|---|---|---|---|---|---|---|
| *HIBCH* | 2 : 191054461-191208919 | 2q32.2 | 3,97e-06 | rs785259 | 0 | NA | NA | 7 | + | 1.742e-09 | NA |
| *AC013468.1* | 2 : 190642909-190643157 | 2q32.2 | 3,97e-06 | rs785259 | 0 | NA | NA | 4 | + | 2.78297982504e-06 | NA |
| *PPP3CA* | 4 : 101944566-102269435 | 4q24 | 4,79e-06 | rs10002927 | 2 | 1.437 | 6 | 0 | NA | NA | NA |
| *ABCC1* | 16 : 16043434-16236931 | 16p13.11 | 4,90e-06 | rs142887759 | 19 | 7.92 | 2b | 0 | NA | NA | Mesenchymal Stem Cell |

**Supplemental Table 3: Genes identified by at least one of the positional, eQTL and chromatin interaction (CI) mappings performed by FUMA based on functional consequences of SNPs with individual p-values ≤ 5 x $10^{-6}$**. Chromosomal positions are based on hg19/*GRCh37* genomic build. For each gene, the lowest GWAS p-value (min GWAS p-value) and the lead SNP (defined as SNP with the lowest p value) in the genomic risk locus to which the gene belongs (locus with at least one SNP with a p-value of association ≤ $10^{-6}$), are indicated. Filled oranges boxes indicate whether the gene was identified by positional, eQTL and/or chromatin interaction (CI) mapping. For positional mapping, the number of SNPs in LD ($r^2$ >0.6) with the lead SNP (nSNPs) as well as the maximum CADD score and minimum RegulomeDB score observed in the gene region are shown. A CADD score >12.37 indicates potential pathogenicity and an RDB score ≤ 4 indicates that the SNP likely lies in a functional location (categories 1 and 2 of RegulomeDB classification identifed as “likely to affect binding”). For eQTL mapping, the number of significant eQTL associations (nSNPs) is reported, together with the direction of effect allele to gene expression for the risk increasing allele of GWAS (eQTL direction) and the associated p-value. For CI mapping, the tissue/cell types for which an interaction was detected are specified. NA : not applicable.

| Gene | Name | p-value | Expressed in human spleen (RPKM) | Malaria OR *Plasmodium* | Red blood cell OR erythrocyte OR erythropoiseis | Spleen | Smooth muscle OR smooth muscular cell | Endothelium OR endothelial cell | Macrophage OR phagocytosis |
|---|---|---|---|---|---|---|---|---|---|
| *CPO* | Carboxypeptidase O | $3.84x10^{-6}$ | 0.291 | | | | | | |
| *CHERP* | Ca2+ homeostasis in endoplasmic reticulum | $6.97x10^{-6}$ | 10.66 | | | | | | |
| *CEBPZ-AS1* | CEBPZ antisense RNA 1 | $1.67x10^{-5}$ | 10.95 | | | | | | |
| *MB* | Myoglobin | $1.68x10^{-5}$ | 0 | Increased levels in severe malaria *Yeo J Infect Dis 2009* | | | | MB plays a role in NO-induced vasodilation and vascular tone regulation *Hendgen-Cotta et al. Trends Cardiovasc Med 2014, Rayner J Biol Chem 2005, and others* | MB scavenges NO, with consequences on the vascular tone *Flögel PNAS 2001* |
| *CTD-3222D15* | CTD | $1.99x10^{-5}$ | ND | | | | | | |
| *IL10RB* | Interleukin 10 receptor subunit β | $3.91x10^{-5}$ | 15.73 | IL-10RB polymorphisms may be protective against severe malaria. *Velavan Immunogenetics 2012* | | | | IL10 improves endothelial function in hypertension. *Kassan Arterioscler Thromb Vasc Biol 2011* | Loss of IL10 signal reduces bacterial killing by macrophages. *Mukhopadhyay J Exp Med 2020; Shouval Gastroenterology 2016 and Immunity 2014.* |

| | | | | | | | | | |
|---|---|---|---|---|---|---|---|---|---|
| *PALLD* | Palladin, cytoskeletal associated protein | $4.44 \times 10^{-5}$ | 21.59 | | PALLD impacts murine erythropoiesis *Liu Blood 2007* | | PALLD plays a role in mesenchymal & muscle cells formation, maturation, migration, and contraction *Ouali BMC Res Notes 2020* | | PALLD regulates phagocytosis . *Sun J Immunol 2017* |
| *SLC35E2* | Solute carrier family 35, member E2 | $4.69 \times 10^{-5}$ | 9.8 | | | | | | |
| *SPARC* | Secreted protein acidic and cysteine rich | $6.68 \times 10^{-5}$ | 123.8 | | SPARC promotes development of erythroid progenitors *Luo et al. Exp Hematol 2012* | SPARC-null mice are splenomegalic and display altered IgM response *Rempel Genes Immun 2007* | SPARC inhibits the proliferation of smooth muscle cells. *Motamed J Cell Biochem 2002* | Splenic sinus endothelial cells express SPARC *Rempel Genes Immun 2007* SPARC impacts the synthesis of collagen IV, present in the basal fibers of splenic sinus walls *Chioran Dev Biol 2017* | Macrophages are a source of SPARC *Hu J Cancer 2020* & *McDonald Am J Physiol Heart Circ Physiol 2018* SPARC induces a M1 polarization *Toba Am J Physiol Cell Physiol 2015* Loss of SPARC promotes macrophage activation in brain tumors *Thomas Brain Pathol 2015* SPARC is produced by macrophages and promotes |

| | | | | | | | | | |
|---|---|---|---|---|---|---|---|---|---|
| | | | | | | | | | metastasis *Sangaletti Cancer Res 2008* But decreases invasiveness in overian cancer *Said Neoplasia 2008* |
| *PDE10A* | Phosphodiesterase 10A | 8.25x10$^{-5}$ | 0.66 | | | PDE10A is involved in spleen-related diving performance *Ilardo Cell 2018* | PDE10A impacts the contraction and proliferation of prostatic and vascular smooth muscle cells. *Hennenberg Prostate 2016, Luo Cardiovasc Res 2021* | | PDE10A plays a role in macrophage-mediated inflammation in the lung *Hsu J Immunol 2021* |

**Supplemental Table 4** : **genes associated with erythrocyte deformability, identified through the MAGMA gene-based association testing**. The p-value for the considered gene is displayed, as well as the transcript abundance as per the NCBI gene database. RPKM : reads per kilobase million. The results of the PubMed literature search addressing the role of each selected gene with relevant keywords (columns) are displayed in red boxes (no association), orange boxes (indirect association) or green boxes (direct associations). Grey boxes : no data available. ND: no data.
